# Additive and epistatic QTL contribute to the adaptation of the fungus *Leptosphaeria maculans* to *Brassica carinata*

**DOI:** 10.64898/2026.07.27.740980

**Authors:** Julie M. Noah, Marie-Hélène Balesdent, Marie Foulongne-Oriol, Mathilde Gorse, Camilla Langlands-Perry, Nicolas Lapalu, Thierry C. Marcel, Benoit Moury, Thierry Rouxel, Jessica L. Soyer

## Abstract

*Leptosphaeria maculans* is a plant-pathogenic fungus that infects *Brassica* species, including *Brassica napus* (oilseed rape). Breeding oilseed rape varieties with genetic resistance is an efficient way to control the disease; however, *L. maculans* can adapt and overcome these resistances. Understanding the mechanisms that enable *L. maculans* to adapt is crucial for managing the emergence of better-adapted isolates. *Brassica carinata*, the Ethiopian Mustard, although closely related to *B. napus*, is considered a nonhost species of *L. maculans* because this fungus cannot infect it. Despite the extreme resistance of *B. carinata*, one natural *L. maculans* isolate has been identified as unable to infect *B. napus*, causing moderate and atypical symptoms on this species. We performed a cross between this isolate and an isolate adapted to *B. napus*, followed by a QTL analysis, which identified seven QTL, each encompassing candidate genes involved in *L. maculans* adaptation to *B. carinata* or *B. napus*. Additionally, we observed transgression in the progeny, wherein a few strains caused significantly more or less aggressive symptoms on both species of *Brassica*. We found that epistasis within the *L. maculans* genome contributes to the observed transgression. These initial findings provide further opportunities to study the adaptive capacities of *L. maculans*, as well as data to initiate analysis of the extreme resistance of *B. carinata* to *L. maculans*.

**Highlights:**

- Seven pathogenicity QTL identified, carrying several interesting candidate genes
- Transgression of some progeny isolates on *B. napus* and *B. carinata* was reported
- Epistasis plays a significant role in the adaptation of *L. maculans* toward host and nonhost *Brassica* species

## Introduction

The prevalence of pathogenic fungi, along with their remarkable adaptive capacities (McDonald and Linde, 2002), poses a serious threat to food safety and human health (Dean *et al*., 2012; Fisher *et al*., 2012). The combination of factors, including global trade, current practices within modern agroecosystems, and the rapid evolution of fungi, represents a high risk to cultivated plants (Fones *et al*., 2020). Unravelling the adaptive mechanisms in pathogenic fungi is thus a highly relevant challenge.

During establishment within their hosts, fungi secrete effectors (proteins, siRNA, or secondary metabolites) that are key in nutrient uptake, fungus development, and/or overcoming of the host immune defence system (Lo Presti *et al*., 2015; Collemare *et al*., 2019). Some effectors, referred to as avirulence proteins (AVR), can be recognized by host resistance proteins. Qualitative host resistance (i.e., monogenic resistance) is widely used in breeding programs to limit the impact of fungal diseases on crops (Nelson *et al*., 2018). Monogenic resistances rely on the gene-for-gene relationship (Flor, 1955) and depend on direct or indirect interaction between the product of a resistance gene (*R*) from the host and the product of an avirulence gene (*AVR*) from the fungus. This interaction can lead either to a complete resistance through a hypersensitive reaction due to local cell death (Balint-Kurti, 2019) or to a partial resistance (e.g., Meile *et al*., 2018; 2023; Jiquel *et al*., 2021; Langlands-Perry *et al*., 2023), both limiting the spread of the pathogen within host tissues. Nonetheless, pathogens may rapidly adapt to plant defences and circumvent monogenic resistance (Brown, 2015; Rouxel and Balesdent, 2017). This presents a significant challenge for plant breeders to develop varieties with resistance that remain effective over a long period. Nonhost resistance (NHR) is a large-spectrum resistance, in which all genotypes of a given plant species are entirely resistant to all genotypes of a given targeted pathogen species (Niks and Marcel, 2009). NHR is based on a combination of defence mechanisms and is, therefore, considered more durable than monogenic host resistance (Fonseca and Mysore, 2019). NHR has been suggested as a promising approach to controlling fungal diseases in agronomy (Ayliffe and Sørensen, 2019; Panstruga and Moscou, 2020).

Factors favouring fungal adaptation are diverse (Sánchez-Vallet *et al*., 2018). Among them, the location of genes in transposable element (TE)-rich regions or dispensable chromosomes contributes to genome evolution (Fudal *et al*., 2009; Faino *et al*., 2016; Hartmann *et al*., 2017; Fokkens *et al*., 2018; Torres *et al*., 2021). Variation of the effector repertoire and reshuffling of genes during sexual reproduction are also important drivers of fungal adaptation (Couch *et al*., 2005; Daverdin *et al*., 2012; Sharma *et al*., 2014; Croll *et al*., 2015; Van Dam *et al*., 2016). These factors may also broaden the host spectrum of pathogens, leading to disease in new plant species (Schulze-Lefert and Panstruga, 2011; Thines, 2019).

The Quantitative Trait-Loci (QTL) and Genome-Wide Association Study (GWAS) approaches, relying on the identification of statistical associations between variations in the genome and in a phenotype within a population, are powerful methods to identify fungal genomic regions, or genes, underlying plant-fungi interaction (Fourie *et al*., 2019; Laurent *et al*., 2021; Louet *et al*., 2021; Dahanayaka and Martin, 2023). For instance, numerous examples of QTL and GWAS analyses have been carried out in the fungus *Zymoseptoria tritici* (Lendenmann *et al*., 2014, 2015, 2016; Zhong *et al*., 2017; Stewart *et al*., 2018; Hartmann *et al*., 2021a; Amezrou *et al*., 2023, 2024; Langlands-Perry *et al*., 2023; Miñana-Posada *et al*., 2025; Stapley *et al*., 2025). To date, few genetic analyses have focused on the genetic determinants underlying host jumps, changes in host range, or adaptation to a nonhost species in fungi.

*Leptosphaeria maculans* is a Dothideomycete fungus that causes stem canker disease of different *Brassica* species and leads to important yield losses in oilseed rape (*Brassica napus*) (West *et al*., 2001; Fitt *et al*., 2008). To date, the favoured method for preventing stem canker disease is to breed resistant oilseed rape cultivars that carry either monogenic resistance (*Rlm* and *LepR* genes) or polygenic, quantitative resistance (Delourme *et al*., 2006; Vasquez-Teuber *et al*., 2024). Although efficient, *Rlm* genes are often not durable, as *L. maculans* displays a significant capacity to adapt and bypass *Rlm* genes deployed in the field (Rouxel *et al*., 2003; Balesdent *et al*., 2022). TE-rich heterochromatin regions are enriched in putative and cloned effector genes, including avirulence genes (*AvrLm*) of *L. maculans* (Rouxel *et al*., 2011; Grandaubert *et al*., 2014; Soyer *et al*., 2014, 2021; Gay *et al*., 2021). In addition, *L. maculans* displays both asexual and sexual reproduction modes with a round of sexual reproduction on oilseed rape residues every year (Rouxel and Balesdent, 2005). Its reproduction regime, together with the location of effector genes in TE-rich regions, favours the emergence of isolates that overcome the genetic resistance of *B. napus* in the field (Rouxel and Balesdent, 2017).

*Brassica carinata*, closely related to *B. napus*, is a nonhost species of *L. maculans* (Noah *et al*., 2024a). However, if NHR is considered more durable than the currently used monogenic resistance, it is not insurmountable (Panstruga and Moscou, 2020). A previous study showed that several field isolates of *L. maculans* were able to cause moderate symptoms on cotyledons of *B. carinata* under controlled conditions, yet unable to cause symptoms on *B. napus*, contrary to other isolates known so far (Noah *et al.,* 2024a). The high adaptive capacity of *L. maculans*, its ability to produce progenies through sexual crosses in controlled conditions, and the availability of a natural isolate causing symptoms on *B. carinata,* make *L. maculans* a perfect model for identifying the genomic regions involved in its adaptation to different plant species through QTL analysis. Here, we use a progeny from the cross between the *B. napus*-adapted reference isolate and an isolate causing moderate symptoms on *B. carinata* (Noah *et al*., 2024a) to identify genomic regions involved in the pathogenicity of *L. maculans* on these two *Brassica* species. Single Nucleotide Polymorphisms (SNPs) were identified in the progeny strains following whole-genome sequencing, and the interaction phenotypes were monitored by inoculating them on several *B. napus* and *B. carinata* genotypes. Our analysis allowed the identification of i) the QTL involved in the pathogenicity of *L. maculans* toward *B. napus* or *B. carinata* and ii) the genetic basis of the transgressions observed in the progeny. Candidate genes in the QTL intervals were identified using previously available genomic and transcriptomic data. This work facilitates a deeper understanding of the mechanisms underlying the adaptation of fungal pathogens to different plant species.

## Materials and Methods

### Fungal material

The *L. maculans* strains used for this study were obtained from a cross between the isolates JN3 and HB10.19 (Noah *et al.,* 2024a), and are hereinafter referred to as the V77.i.j progeny strains. Of the 102 progeny strains, 10 did not sporulate and were therefore unable to be used for the pathogenicity assays. Fungal strains were stored in the long term at 4°C on 1% malt-2% agar slant tubes, and pycnidiospore suspensions were produced as described in Ansan-Melayah *et al*. (1995) and stored at - 20°C.

### Plant genotypes

We inoculated five genotypes of *B. carinata* (lines D5.6.12, BRA.2264, BRA.2452, BRA.1029, and BRA.1043; Noah *et al.,* 2024a; IPK Genebank https://www.ipk-gatersleben.de/en/research/genebank) and three genotypes of *B. napus* (cvs. Yudal carrying resistance gene *RlmSTEE98*, ES-Astrid carrying resistance gene *Rlm9*, and Westar with no known resistance genes) to assess the infection phenotypes of the V77 progeny and parental isolates.

### Pathogenicity assays

Pathogenicity assays at the cotyledon stage were performed as described in Balesdent *et al*. (2022) on 12-day-old seedlings of *B. napus* and *B. carinata*, with three independent experiments each including 12 technical replicates (i.e., 12 inoculation points for each *L. maculans* isolate-plant genotype combination). We inoculated plants of *B. carinata* and *B. napus* with pycnidiospore suspensions (10^7^ pycnidiospores.ml^-1^) of JN3, HB10.19, and their 92 progeny strains.

### Phenotypic traits evaluated and statistical analyses

To identify the genetic determinants involved in the pathogenicity of *L. maculans* towards *B. carinata* and *B. napus*, we evaluated three different phenotypic traits: i) the average disease score using the LeMaSQ scale (Noah *et al*., 2024b) ranging from 1 (non-adapted isolate) to 9 (adapted isolate), as the symptoms caused by the HB10.19 isolate and the V77 progeny strains on *B. napus* and *B. carinata* cotyledons were not represented in the IMASCORE rating scale (Balesdent *et al*., 2001); ii) the virulence percentage, corresponding to the ratio between the number of virulent inoculation points (i.e., symptom score from 7 to 9) and the total number of inoculation points evaluated per isolate; iii) the symptom area, measured with the ImageJ software (Schneider *et al*., 2012). These phenotypic traits were measured 12 days post-inoculation (dpi).

To identify possible significant differences between experiments and to estimate the effects of fungal strains and replicates, we performed Wilcoxon-Mann-Whitney tests, Kruskal-Wallis tests, and ANOVAs using RStudio version 2023.09.0+463 (RStudio Team, 2020) with the R software version 4.3.1 (R Core Team, 2019). Broad-sense heritability (*H*²) is the proportion of total phenotypic variance (VP) that is due to total genetic variance (VG). This includes all genetic effects, such as additive, dominance, and epistasis (Falconer and Mackay, 1996). We calculated the heritability *H²* as described in Langlands-Perry *et al*. (2021) using the formula *H²* = σg2/(σg2+ σe2), where σg2 is the genotypic variance and σe2 the residual variance.

### DNA extraction, whole-genome sequencing, and variant calling

Fungal strains were maintained on V8-juice agar medium (Ansan-Melayah *et al*., 1995), grown at 25°C for five to seven days before harvest as described in Soyer *et al*. (2021), and lyophilized for 24 hours before DNA extraction. Extraction of the genomic DNA from the parental isolates JN3 and HB10.19 and the 92 V77 progeny strains was realized using 20 to 30 mg lyophilized mycelium using the DNeasy Plant Mini Kit (QIAGEN S.A., Courtaboeuf, France) as described by Jacques *et al*. (2021). The DNA of the parental isolates and progeny strains was sequenced by Genewiz, Inc. (Genewiz NGS Laboratory, Takeley, UK) using 2x100 bp reads for the parental isolates, and 2x150 bp reads for the progeny strains on a HiSeq-2000 sequencing system. The variant calling was performed as described in Amezrou *et al*. (2023) by mapping the reads onto the reference genome of *L. maculans* isolate JN3 (Dutreux *et al*., 2018).

### Genetic linkage map

To construct the linkage map of the JN3 x HB10.19 cross, we used a file containing all markers (SNPs) from the progeny strains as input for the Multipoint ultra-dense software (MultiQTL Ltd., Haifa, Israel; Mester *et al*., 2003) as described in Langlands-Perry *et al*. (2021). All SNP with more than 5.43% missing data were filtered out (i.e., the marker was absent in a maximum of 5 out of 92 progeny strains), and the maximum Chi-Square value to filter markers with distorted segregation was set to 2.6. The Multipoint ultra-dense software established linkage groups (LG) based on genetically-linked markers. Co-segregating markers were grouped in bins using a "twin algorithm" (Ronin *et al*., 2017). We chose a skeleton marker with a minimal rate of missing data to represent the entire set of markers within each bin. The ordering of skeleton markers follows a Guided Evolutionary Strategy heuristic based on the minimum distance between adjacent skeleton markers to minimize the total length of the genetic map (Mester *et al*., 2003). As there was too much missing data for markers on the dispensable chromosome of *L. maculans*, we increased the threshold percentage of missing data to 10% for the LG corresponding to this chromosome. The genetic map generated by Multipoint with the skeleton markers was visualized using MapChart (Voorrips, 2002).

### QTL analyses

QTL mapping was performed as described by Langlands-Perry *et al*. (2021) using RStudio with the R/qtl package (Broman *et al*., 2003; Arends *et al*., 2010). QTL confidence intervals were determined using the LOD support, with a drop of 1 LOD score value and the "expandtomarkers" argument set as “true”. In total, we analysed 72 datasets (three traits for three experiments performed on eight *Brassica* genotypes). We considered QTL with *p*-values <0.05 or 0.1 (LOD score threshold) to be relevant, and we selected them for the following analyses. For QTL identified through multiple experimental replicates or multiple phenotypic traits (≥2), we considered the most frequent flanking SNP to define confidence intervals. When multiple QTL were identified in a dataset, we tested the occurrence of an epistatic interaction between them using the R/qtl package as described by Langlands-Perry *et al*. (2021). We used Chi-square tests (*p*-values<0.05) to evaluate the significant enrichment in different predicted genomic features (effector genes, CAZyme-encoding genes, TE-associated genes, or genes located within a heterochromatin region; Dutreux *et al.,* 2018; Soyer *et al.,* 2021) within the QTL intervals (RStudio Team, 2020).

### Analysis of phenotypic transgressions

In the *L. maculans* V77 progeny, some strains exhibited higher or lower pathogenicity values (i.e., disease score) than the two parental isolates on *B. carinata* and *B. napus* cotyledons (including 15 isolates showing a significantly higher pathogenicity value; Noah *et al.,* 2024a). To investigate the putative transgressive status of these strains, Wilcoxon-Mann-Whitney tests with a *p-*value threshold of 0.05 were performed to test whether their disease scores on *B. carinata* or *B. napus* were significantly different from those of the parental isolates. To investigate the genetic basis of the observed phenotypic transgressions, we examined epistatic relationships between pairs of markers in the entire *L. maculans* genome using ANOVAs performed in RStudio (with the ANOVA model: disease score ∼ marker 1*marker 2). We used a Bonferroni correction to adjust for an overall *p*-value of 0.05, correcting for multiple ANOVA tests. Using too many markers would result in a large number of possible marker pairs, thereby leading to a substantial number of ANOVA tests and an increased risk of low statistical power. Thus, we selected 194 molecular markers located at an average interval of 8.6 cM on the genetic map (i.e., the total length of the genetic map divided by the number of markers) to perform the ANOVA, including the peak markers of the previously identified QTL intervals. We kept the markers that were conserved across the genotypic data of the progeny strains. When an epistatic relationship was found between two markers, we performed a Fisher’s exact test to determine whether, for this pair of markers, the transgressive strains carried more frequently the favourable (i.e., higher pathogenicity) allelic combination(s) than the non-transgressive strains.

### Identification of candidate genes within QTL intervals

Markers flanking each QTL were used to identify the genomic intervals on the reference genome of the JN3 *L. maculans* isolate (Dutreux *et al.,* 2018) corresponding to the QTL confidence intervals. Genes of *L. maculans* annotated within these intervals were listed and screened for differential expression during *B. napus* infection (data from Gay *et al.,* 2021) and for their location in a euchromatin or heterochromatin domain during axenic culture (data from Soyer *et al.,* 2021). We aligned Illumina reads of the HB10.19 isolate with the reference genome of *L. maculans* to assess nucleotide sequence variations between the parental isolates for the genes included in the QTL intervals. For each sequence polymorphism between HB10.19 and JN3, we predicted the type of modification (synonymous or non-synonymous) on the encoded protein using SnpEff v5.1 (Cingolani *et al*., 2012).

## Results

### Generation of an ultra-dense genetic map of *Leptosphaeria maculans*

The complete genome of the 92 V77 progeny strains from the cross between JN3 and HB10.19 was sequenced using an Illumina strategy; 46,585 SNPs were identified in the progeny and used as input in the Multipoint ultra-dense software. After filtering for missing data, we retained 35,613 markers for linkage analysis. The genetic map comprised 1,466 skeleton markers and 34,147 co-segregating markers (i.e., markers sharing the same genetic position as a skeleton marker; Table 1). The average distance between skeleton markers was 1.19 cM, corresponding to 29,246 bp and representing an average map density of 21 cM/Mb. The genetic map comprised 20 LG (LG 0 to LG 19, corresponding to SuperContigs (SC) 0 to 19 of the genome of *L. maculans*) for a total size of 1,721 cM, covering 42.3 Mb (92.1%) of the *L. maculans* reference genome (Dutreux *et al*., 2018; Fig. 1). Both parental isolates carry the *L. maculans* dispensable chromosome, corresponding to SC 19 (i.e., LG 19). However, 12 out of the 92 progeny strains lack the dispensable chromosome, and markers associated with SC 19 were absent in these strains. The complete genetic map (Fig. 1) displayed large regions not covered by molecular markers due to TE, as we chose to exclude the SNPs identified inside the TE-rich regions. Moreover, we identified a deletion of a region encompassing almost 26 kb on the LG 3, located in an AT-isochore, as markers corresponding to this region were missing in the genomes of 44 out of the 92 progeny isolates, while the markers were present in the parental isolates. Based on the relationship between the genetic markers and their physical positions along the genome, we did not detect any chromosomal rearrangement among the progeny strains compared to the parental isolates (Fig. S1). We identified 4,096 crossovers (CO) in the progeny strains, representing an average number of 43 CO per strain and an average of 2.18 CO per linkage group per strain (Table S1). The number of CO per LG was correlated with the size of the LG (Pearson’s *r*= 0.94, *p*-value<0.05), with the smallest LG, LG 19, having the lowest number of CO. For the core chromosomes, the density of CO was not correlated with the size of the LG (i.e., crossover number divided by the physical size of each LG). Nevertheless, the dispensable chromosome (LG 19) of *L. maculans* had the lowest CO density (CO density on LG 19 = 0.008/kb; LG 0 – LG 18 CO density = 0.1/kb). The lowest CO density identified within the dispensable chromosome of *L. maculans* might be associated with the fact that the dispensable chromosome was absent or partially conserved in some progeny strain (12 progeny strains did not have molecular markers within LG 19, three had very few markers (less than ten), 26 had large parts of that linkage group without markers and only six had recombination in this linkage group).

**Fig. 1.**
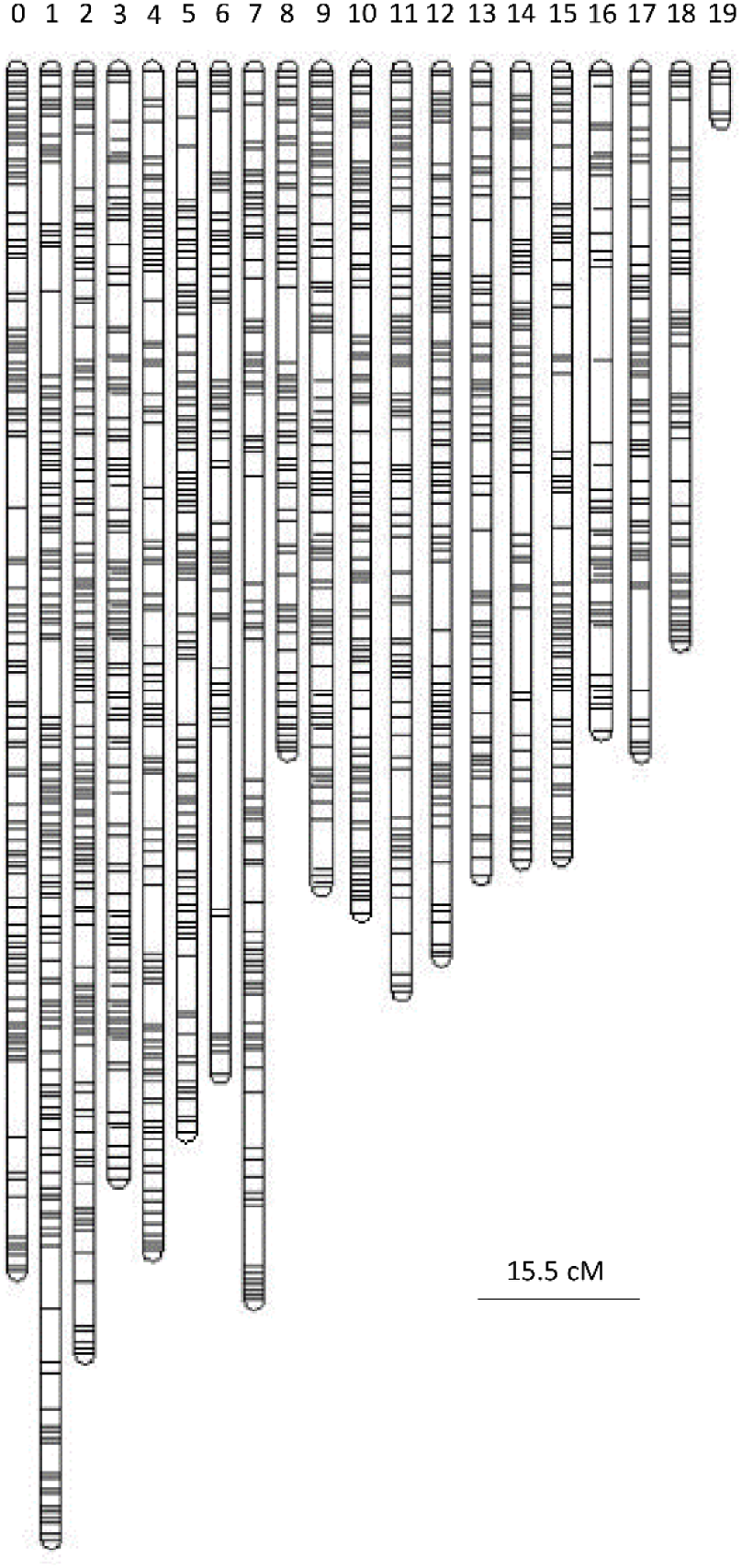
*Leptosphaeria maculans* genetic map determined following the cross between isolates JN3 and HB10.19.

**Table 1.** Characteristics of the genetic linkage map of the cross between *Leptosphaeria maculans* isolates JN3 and HB10.19.

| Linkage Group<sup>a</sup> | Number of markers | Number of skeleton markers<sup>b</sup> | Average number markers/bin<sup>b</sup> | Total genetic distance (cM) | Average distance between skeleton markers (cM) | Total physical distance (kb)<sup>c</sup> |
| --- | --- | --- | --- | --- | --- | --- |
| 0 | 2,397 | 102 | 23.50 | 114.61 | 1.12 | 3,606 |
| 1 | 2,876 | 127 | 22.65 | 140.21 | 1.10 | 3,546 |
| 2 | 3,149 | 119 | 26.46 | 122.51 | 1.03 | 3,418 |
| 3 | 2,885 | 98 | 29.44 | 105.83 | 1.08 | 2,905 |
| 4 | 2,017 | 90 | 22.41 | 112.73 | 1.25 | 2,871 |
| 5 | 2,277 | 93 | 24.48 | 101.39 | 1.09 | 2,518 |
| 6 | 1,756 | 58 | 30.28 | 95.81 | 1.65 | 2,528 |
| 7 | 1,721 | 79 | 21.78 | 117.42 | 1.49 | 2,359 |
| 8 | 1,627 | 61 | 26.67 | 65.02 | 1.07 | 2,067 |
| 9 | 1,962 | 77 | 25.48 | 77.88 | 1.01 | 2,096 |
| 10 | 2,095 | 80 | 26.19 | 80.40 | 1.01 | 1,865 |
| 11 | 1,843 | 75 | 24.57 | 87.86 | 1.17 | 1,783 |
| 12 | 1,706 | 78 | 21.87 | 84.60 | 1.08 | 1,788 |
| 13 | 1,473 | 59 | 24.97 | 76.80 | 1.30 | 1,636 |
| 14 | 1,397 | 60 | 23.28 | 75.35 | 1.26 | 1,471 |
| 15 | 1,113 | 62 | 17.95 | 75.11 | 1.21 | 1,437 |
| 16 | 948 | 42 | 22.57 | 63.12 | 1.50 | 1,330 |
| 17 | 1,108 | 54 | 20.52 | 65.17 | 1.21 | 1,161 |
| 18 | 1,144 | 47 | 24.34 | 54.60 | 1.16 | 1,187 |
| 19 | 119 | 5 | 23.80 | 4.67 | 0.93 | 773 |
| Total/Mean | 35,613 | 1,466 | 24.29 | 1,721.09 | 1.19 | 42,345 |

<sup>a</sup>Linkage group represents the chromosomes (i.e. Super-Contigs) of *L. maculans*; <sup>b</sup>A bin is composed of a skeleton marker and the linked markers that segregate with this skeleton marker in the progeny; <sup>c</sup>according to the reference genome of *L. maculans* isolate JN3 (Dutreux *et al.*, 2018).

The genetic map was generated using single nucleotide polymorphisms identified in the progeny strains from the cross between the *Leptosphaeria maculans* isolates JN3 and HB10.19 (Materials and Methods). The numbers represent the linkage groups (LG) of the genetic map and correspond to the chromosome numbers (i.e. Super-Contigs; Dutreux *et al*., 2018). Each bar line within the LG represents the position of a bin (one skeleton marker associated with its linked markers). The visualization of the genetic map was obtained using MapChart.

### Continuum of aggressiveness of the progeny strains on *Brassica napus* and *Brassica carinata* cotyledons

In a previous study, we analysed the pathogenicity of several *L. maculans* isolates, including HB10.19, moderately adapted to *B. carinata*, and JN3, adapted to *B. napus*, on 52 genotypes of (Noah *et al.,* 2024a). Based on these pathogenicity assays, we selected five *B. carinata* genotypes displaying different levels of resistance or susceptibility toward HB10.19 and V77.1.11, a progeny strain exhibiting a positive transgressive phenotype on these *B. carinata* genotypes (i.e., having a disease score superior to that of HB10.19; Noah *et al.,* 2024a; Fig. 2, Fig. S2, and S3). The three *B. napus* genotypes, Yudal, Westar, and ES-Astrid, were susceptible to the isolates JN3 and V77.1.11, and were used in this study (Fig. 2, and Fig. S4). To identify the genomic regions involved in the pathogenicity of *L. maculans* towards *B. carinata* and *B. napus*, we analysed the phenotypes of the progeny of the JN3xHB10.19 cross on eight *B. carinata* and *B. napus* genotypes. We combined the three independent experiments to calculate the broad-sense heritability (*H²*) of the three phenotypic traits assessed (i.e., the disease score, symptom area, and virulence percentage). All the genotypes of both *Brassica* spp. except for *B. napus* cv. Westar, exhibited high (≥ 55.6%) broad-sense heritability values for the phenotypic traits (Table 2), supporting that the variability of these traits is primarily due to the genetic differences between progeny individuals. In the *L. maculans* progeny, the phenotypic traits were positively correlated between all pairs of *Brassica* spp. genotypes. However, correlations were stronger within lines of *B. carinata* or *B. napus* than between *B. carinata* and *B. napus* lines (Fig. S5). Higher positive correlations were observed between the virulence percentage and the symptom area than between the disease score on both *Brassica* species (Fig. S5). Moreover, the three phenotypic traits positively correlated for each *Brassica* genotype (Fig. S6 and S7), suggesting that at least some of the genetic determinants are shared for these traits. Pathogenicity of the parental isolates JN3 and HB10.19 on all the *B. napus* or *B. carinata* genotypes tested here behaved as already demonstrated (Noah *et al*., 2024a), with JN3 causing typical greyish symptoms on *B. napus*, but causing a localized black symptom at the inoculation point on the *B. carinata* genotypes, while HB10.19 caused moderate symptoms (i.e., greyish symptoms with a black margin on *B. carinata* and a localized black symptom on *B. napus*). The progeny and parental isolates displayed a continuum of aggressiveness on all genotypes of *B. carinata* and *B. napus* (Fig. 2, Fig. S3-S4 and S8-S9; Table S2). Twenty-five progeny strains could not cause a disease score higher than three on all *Brassica* genotypes, showing that a significant part of the progeny could not cause symptoms typical of a disease on the *Brassica* genotypes inoculated. Five progeny strains displayed a disease score ≥6 (symptoms typical of a pathogenic phenotype, with greyish lesions not bordered by a black margin) on the three *B. napus* genotypes, suggesting an adaptation of these strains to the plant (Fig. 2, S4 and S8; Table S2). For *B. carinata*, between two and 12 progeny strains were capable of inducing typical disease symptoms depending on the inoculated genotype, suggesting variable responses among *B. carinata* and *L. maculans* genotypes to the infection (Fig. 2, S3, and S9; Table S2). Only four progeny strains induced disease symptoms on all *Brassica* species, indicating that the combination of parental abilities to infect both *B. napus* and *B. carinata* is possible but rare among the progeny. The continuous variation of the phenotypic traits assessed in the progeny on different *B. napus* and *B. carinata* genotypes suggests that the pathogenicity of *L. maculans* on both *Brassica* species is under polygenic inheritance.

**Fig. 2.**
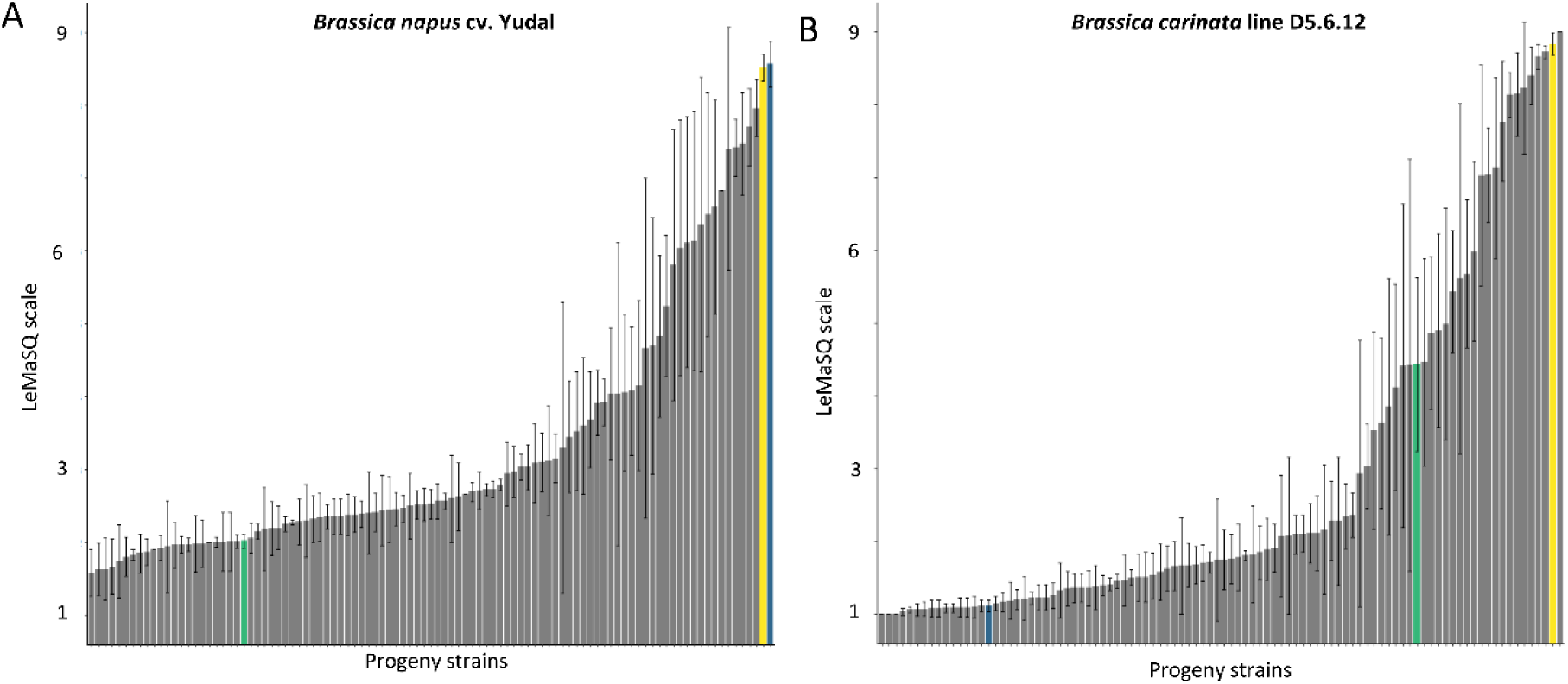
Distribution of disease scores caused by the *Leptosphaeria maculans* V77 progeny strains on A) *Brassica napus* and B) *Brassica carinata* cotyledons. Symptoms were evaluated following the LeMaSQ scale. Dark blue bar, isolate JN3; green bar, isolate HB10.19; grey bars, V77 progeny strains; yellow, strain V77.1.11. Thin bars represent the standard deviation for each strain.

**Fig. 3.**
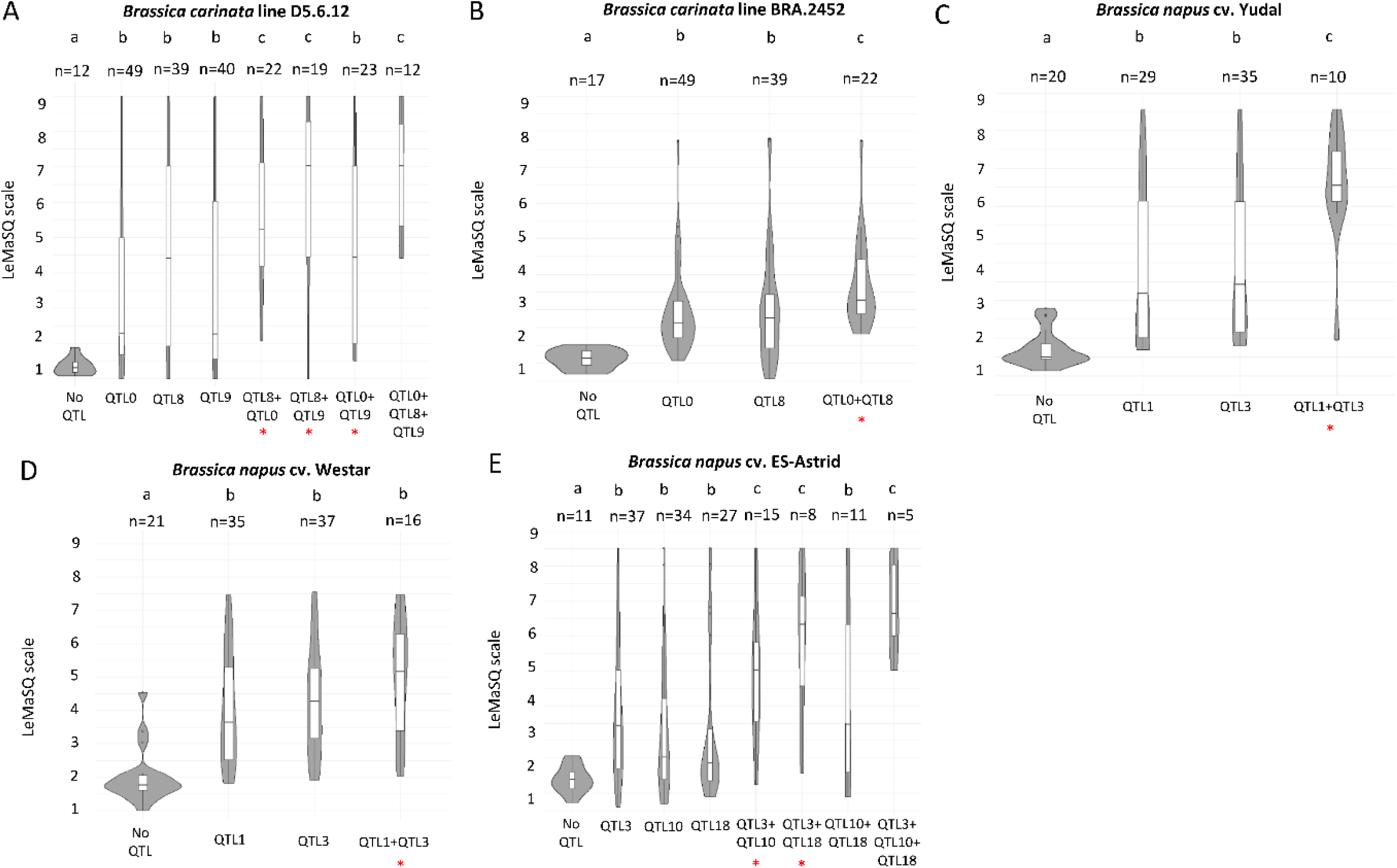
Distribution of phenotypes corresponding to the different allelic combinations at the QTL identified in the V77 progeny. Letters “a”, “b”, and “c” indicate allelic combinations corresponding to significantly different average disease scores (Wilcoxon test; *p*-value<0.05).“QTLx” indicates a favourable allele at the QTL identified on linkage group x. “No QTL”, strains carrying no favourable alleles at the QTL on linkage group x. “n=x” indicates the number of strains in each group. The red star indicates an epistasis between the QTL.

**Table 2.** Broad-sense heritability (*H²*) of the phenotypic traits evaluated for the pathogenicity of Leptosphaeria maculans *towards* Brassica napus *and* Brassica carinata.

| Species | Genotype | $H^2$ (%) | | |
| --- | --- | --- | --- | --- |
|  |  | Disease score | Symptom area | Virulence percentage |
| *Brassica napus* | ES-Astrid | 79.3 | 70.7 | 76.5 |
|  | Yudal | 67.2 | 55.6 | 55.6 |
|  | Westar | 52.6 | 45.1 | 27.9 |
| *Brassica carinata* | D5.6.12 | 82.3 | 68.4 | 71.9 |
|  | BRA.2264 | 83.6 | 69.5 | 62.2 |
|  | BRA.2452 | 80 | 68.4 | 69.1 |
|  | BRA.1029 | 74 | 70.8 | 80.3 |
|  | BRA.1043 | 82.9 | 70.7 | 64 |

### Cultivar or species-specific QTL underlie the pathogenicity of *Leptosphaeria maculans* toward *Brassica napus* or *Brassica carinata*

We mapped the QTL corresponding to the phenotypic traits assessed on the *B. carinata* and *B. napus* genotypes. As the results for interactions between the progeny and the *Brassica* genotypes showed significant differences among the three experimental replicates (ANOVA, *p*-value>0.05), we analysed the results from each replicate separately. All QTL intervals identified overlapped across the individual replicates (Tables S3 and S4). The three phenotypic traits measured led to the identification of the same QTL (Table S3 and S4), consistent with the positive correlation observed among the phenotypes (Fig. S6 and S7). Thus, we describe here only the QTL identified with the “disease score” trait, which explained the highest percentage of phenotypic variance (QTL identified for all phenotypic traits are detailed in Tables S3 and S4). A QTL was considered robust when identified in at least two experimental replicates (regardless of the phenotypic trait for which the QTL was identified), and QTL intervals were defined by selecting the flanking markers that were the most identified among all replicates. Three and four QTL were identified for the pathogenicity towards *B. carinata* and *B. napus*, respectively (Fig. S10 and Table 3), and were located on seven different linkage groups. The QTL were located on LG 1, 3, 10, and 18 for pathogenicity towards *B. napus* (named hereafter QTL_LG1, QTL_LG3, QTL_LG10, and QTL_LG18) and on LG 0, 8, and 9 for pathogenicity on *B. carinata* (hereafter referred to as QTL_LG0, QTL_LG8, and QTL_LG9). Each of the four QTL for pathogenicity of *L. maculans* on *B. napus* explained 5 to 29% of the phenotypic variation (Table S3), with the QTL identified on LG 3, contributing to the highest variation for all genotypes of *B. napus*. The three QTL for pathogenicity on *B. carinata* explained between 13.5% and 67.6% of the phenotypic variation (Table S4), with QTL_LG8, identified across all inoculated *B. carinata* genotypes, contributing the most to the variation.

**Table 3.** QTL involved in the pathogenicity of *Leptosphaeria maculans* towards *Brassica napus* and *Brassica carinata*.

| Linkage group | Species | Cultivar /Line | LOD score<sup>a</sup> | Confidence interval (cM) | Confidence interval (pb) | Size (kb) | % phenotypic variation explained<sup>b</sup> | Parental allele conferring higher disease scores | Number of genes in the QTL interval |
| --- | --- | --- | --- | --- | --- | --- | --- | --- | --- |
| 1 | *Brassica napus* | Yudal, Westar | 1.2 to 7.3 | 47.38 - 64.48 | 1,037,697 – 1,373,089 | 335 | 5 to 19.9 | JN3 | 135 |
| 3 |  | ES-Astrid, Yudal, Westar | 1.8 to 8 | 0 - 5.04 | 200,085 – 341,458 | 141 | 8.5 to 28.9 | JN3 | 27 |
| 10 |  | ES-Astrid | 2.9 to 4 | 67.95 - 76.12 | 1,542,222 – 1,792,971 | 250 | 11.1 to 11.3 | JN3 | 76 |
| 18 |  | ES-Astrid | 2.2 to 3.6 | 25.16 - 53 | 760,523 – 896,682 | 136 | 8.2 to 8.7 | JN3 | 64 |
| 0 | *Brassica carinata* | D5.6.12<br>BRA.2452 | 2.8 to 13 | 41.75 - 49.33 | 1,516,732 – 1,838,035 | 321 | 16.9 to 41.5 | HB10.19 | 97 |
| 8 |  | D5.6.12<br>BRA.2264<br>BRA.2452<br>BRA.1029<br>BRA.1043 | 2.7 to 22.5 | 63.43 - 65.02 | 1,451,266 – 2,045,345 | 594 | 13.5 to 67.6 | HB10.19 | 28 |
| 9 |  | D5.6.12 | 2.9 to 5.6 | 23.37 - 33.54 | 923,181 – 1,014,686 | 91 | 16.6 to 17.2 | HB10.19 | 36 |

We present only the QTL identified using the disease score phenotype, as the symptom area and virulence frequency traits allowed the detection of the same QTL. Results correspond to the combination of the three independent experiments.
<sup>a</sup>The LOD score *p-value* threshold is 0.05 or 0.1 (Tables S3 and S4);
<sup>b</sup>Proportion of phenotypic variation attributed to each of the QTL identified using the disease score and for the three experimental replicates.

None of these QTL were conserved between the pathogenicity of *L. maculans* on *B. napus* and *B. carinata* species. For each species, we identified one QTL involved in the pathogenicity of *L. maculans* towards all the genotypes of *B. napus* (QTL_LG3) or *B. carinata* (QTL_LG8), suggesting species specificity of these QTL, whereas other QTL were genotype-specific (Table 3).

### Progeny strains carrying several favourable alleles at the QTL are more aggressive towards *Brassica napus* or *Brassica carinata*

We assessed whether progeny strains combining in their genome favourable alleles of several QTL identified on *B. napus* or *B. carinata* were more pathogenic than strains carrying only one favourable allele of one of those QTL. The favourable alleles at the QTL identified on *B. carinata* lines D5.6.12 and BRA.2452 had a positive additive effect on the disease score (Fig. 3A, B). Progeny strains having the favourable alleles at the QTL_LG8 and QTL_0, or QTL_LG8 and QTL_LG9 involved in the pathogenicity of *L. maculans* towards *B. carinata* line D5.6.12 were more aggressive on this line than strains carrying only one QTL (either QTL_LG0, QTL_LG8, or QTL_LG9). However, these progeny strains were similarly aggressive as progeny strains carrying the favourable alleles of the three QTL (Fig. 3A), indicating that QTL_LG9 and QTL_LG0 contributed little to the observed phenotype on *B. carinata* line D5.6.12. Altogether, these results suggest that the genes embedded in the QTL_LG8 are very important for the pathogenicity of *L. maculans* on *B. carinata* as this QTL displayed a higher effect on the observed phenotype in combination with another QTL compared to the QTL_LG0 and QTL_LG9.

QTL_LG1 and QTL_LG3, involved in pathogenicity on *B. napus* cv. Yudal had an additive effect (Fig. 3C). This additive effect was not observed on the cv. Westar (Fig. 3D), suggesting that the combination of favourable alleles at QTL_LG1 and QTL_LG3 is not required for higher pathogenicity on this cultivar compared to cv. Yudal. An additive effect was also detected for some pathogenicity QTL identified on *B. napus* cv. ES-Astrid (Fig. 3E). The presence of the favourable allele of QTL_LG3 in combination with the favourable allele of QTL_LG10 or QTL_LG18 in the progeny strains led to a higher pathogenicity of the strains on cv. ES-Astrid compared to favourable alleles of QTL_LG3 alone (Fig. 3E). In addition, these combinations had the same effect on *L. maculans* pathogenicity, suggesting QTL_LG3 is the main contributor to successful infection on *B. napus* cv. ES-Astrid. These results indicate that the additive effect of a QTL enhances *L. maculans* pathogenicity on both *B. napus* and *B. carinata*, with a single QTL making a major contribution in both cases.

### Epistasis between QTLs contributes to pathogenicity

In addition to the additive effects of multiple QTL in *L. maculans*, we investigated epistasis (i.e., interactions between two or more genes or alleles at the genome-wide level) in the pathogenicity of the fungus toward *B. napus* and *B. carinata*. On *B. carinata* line D5.6.12 and BRA.2452, we detected significant epistatic effects between QTL_LG0 and QTL_LG8 (explaining from 6.9 to 18.5% of the phenotypic variation), between QTL_LG0 and QTL_LG9 (3-6%), and between QTL_LG8 and QTL_LG9 (6.4 to 16.2%) (Table 4). On *B. napus* cvs. Yudal and Westar, we observed significant epistasis between QTL_LG1 and QTL_LG3 (3-8% of the explained phenotypic variation; Table 4). On *B. napus* cv. ES-Astrid, the epistatic interaction between QTL_LG3 and QTL_LG18 explained up to 20% of the phenotypic variation, whereas the epistatic interaction between QTL_LG3 and QTL_LG10 explained up to 11% of the phenotypic variation. All detected epistases were positive and synergistic: the aggressiveness of *L. maculans* strains carrying two favourable QTL alleles (i.e., alleles that enhanced aggressiveness compared to the less aggressive parental strain) was higher than expected by a purely additive effect between the QTL alleles (Fig. 3). These results indicate that the pathogenicity QTL identified toward *B. napus* and *B. carinata* exhibit epistasis, contributing to the overall explained phenotypic variance.

**Table 4.** Summary of epistatic interactions between the additive QTL involved in the pathogenicity of *Leptosphaeria maculans* towards *Brassica napus* or *Brassica carinata*.

| Species | Cultivars | QTL pair | Epistasis<sup>a</sup> | p-value |
| --- | --- | --- | --- | --- |
| *Brassica carinata* | D5.6.12, BRA.2452 | QTL_LG0 and QTL_LG8 | 6.9 to 18.5 | < 0.001 |
|  | D5.6.12 | QTL_LG0 and QTL_LG9 | 3 to 6 | < 0.05 |
|  | D5.6.12 | QTL_LG8 and QTL_LG9 | 6.4 to 16.2 | < 0.001 |
| *Brassica napus* | ES-Astrid | QTL_LG3 and QTL_LG10 | 4.6 to 11 | < 0.01 |
|  | ES-Astrid | QTL_LG3 and QTL_LG18 | 5.4 to 20 | < 0.01 |
|  | Yudal, Westar | QTL_LG1 and QTL_LG3 | 3 to 8.1 | < 0.01 |

All detected epistatic interactions were positive.
<sup>a</sup>Range of values obtained for the different independent experiments and phenotypic traits explained. QTL\_LGx, indicates QTL identified on linkage group (LG) x.

### Epistasis between markers in the *Leptosphaeria maculans* genome contributes to the transgression observed in the progeny strains

The V77 progeny strains showed both positive and negative transgressive disease score phenotypes on all *B. napus* and *B. carinata* genotypes, indicating they cause significantly more or fewer symptoms than the two parental isolates (Fig. 2, Fig. S3, S4, S8, and S9, and Table S2). Depending on the considered *B. carinata* genotype, there were between four and 16 progeny strains showing significant positive transgression, and between zero and 21 progeny strains showing significant negative transgression (Wilcoxon test, *p-*value < 0.05). On the other hand, between three and six progeny strains showed significant negative transgression on all *B. napus,* whereas no progeny strains showed significant positive transgression on this plant species (Wilcoxon test, *p-*value < 0.05). To decipher the genetic bases of the phenotypic transgressions observed in the *L. maculans* progeny, we analysed the involvement of the epistasis between pairs of SNPs distributed over the entire genome in the transgression. The ANOVA tests revealed no epistasis with a significant impact on the phenotypes assessed on *B. napus* (corrected *p-*value > 0.05). In contrast, significant epistatic interactions were detected for four pairs of markers (M_27480*M_01498, M_27480*M_01669, M_11990*M_28286, and M_23071*M_42047) for the phenotype on *B. carinata* lines D5.6.12, BRA.1043, and BRA.2264 (corrected *p-*value < 0.05) (Fig. 4; Table 5).

**Fig. 4.**
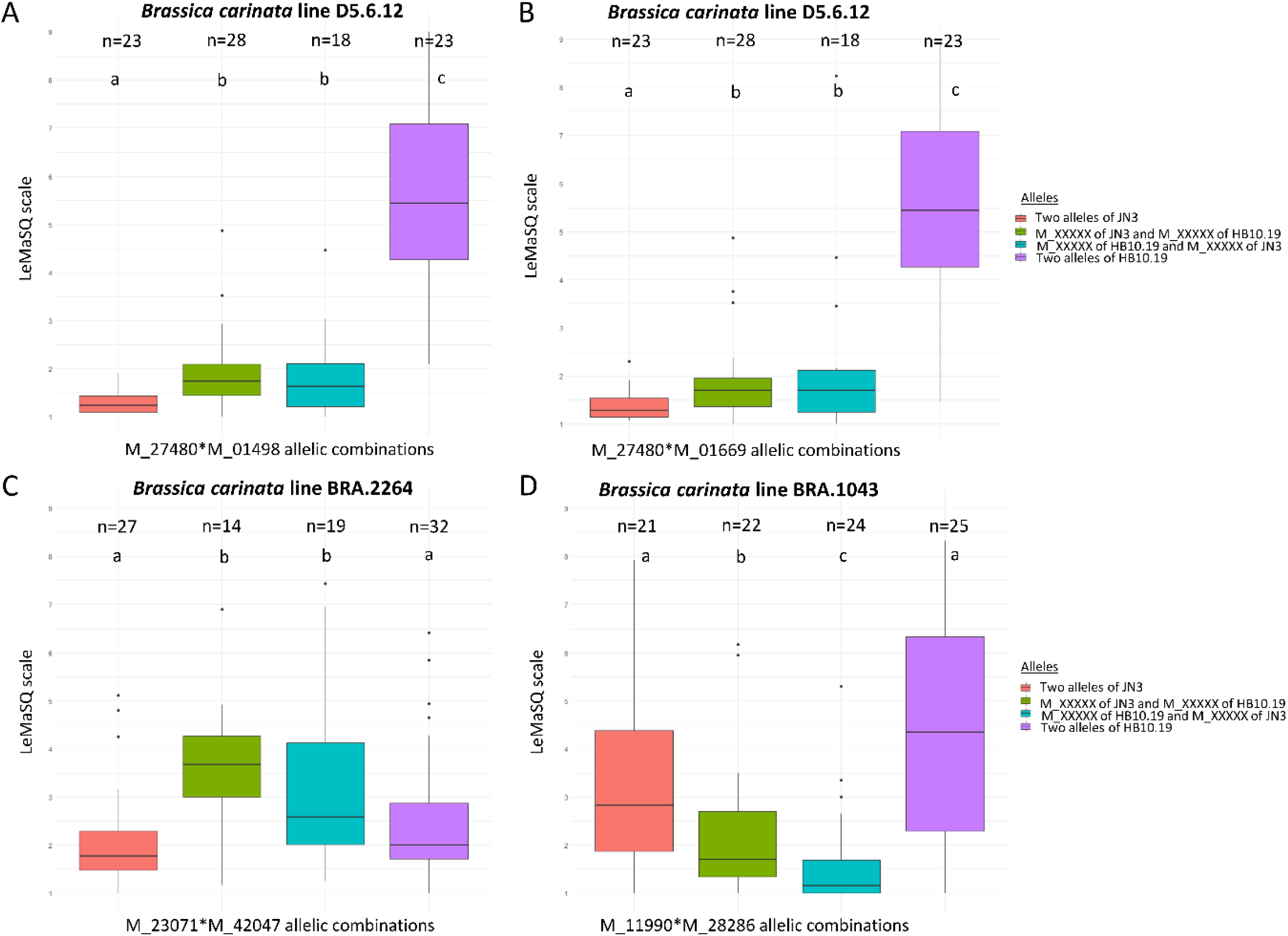
Significant epistatic effects detected for pairs of markers determining pathogenicity (disease score) of the *Leptosphaeria maculans* progeny in *Brassica carinata*. A) Disease score of the progeny on *B. carinata* line D5.6.12 for the four allelic classes of markers M_27480 and M_01498, B) Disease score of the progeny on *B. carinata* line D5.6.12 for the four allelic classes of markers M_27480 and M_01669, C) Disease score of the progeny on *B. carinata* line BRA.2264 for the four allelic classes of markers M_23071 and M_42047, D) Disease score of the progeny on *B. carinata* line BRA.1049 for the four allelic classes of markers M_11990 and M_28286. n=X indicates the number of progeny strains per group of allelic combination. Letters “a”, “b” and “c” indicate allelic combinations corresponding to significantly different average disease scores (Wilcoxon test; *p*-value<0.05).

**Table 5.**
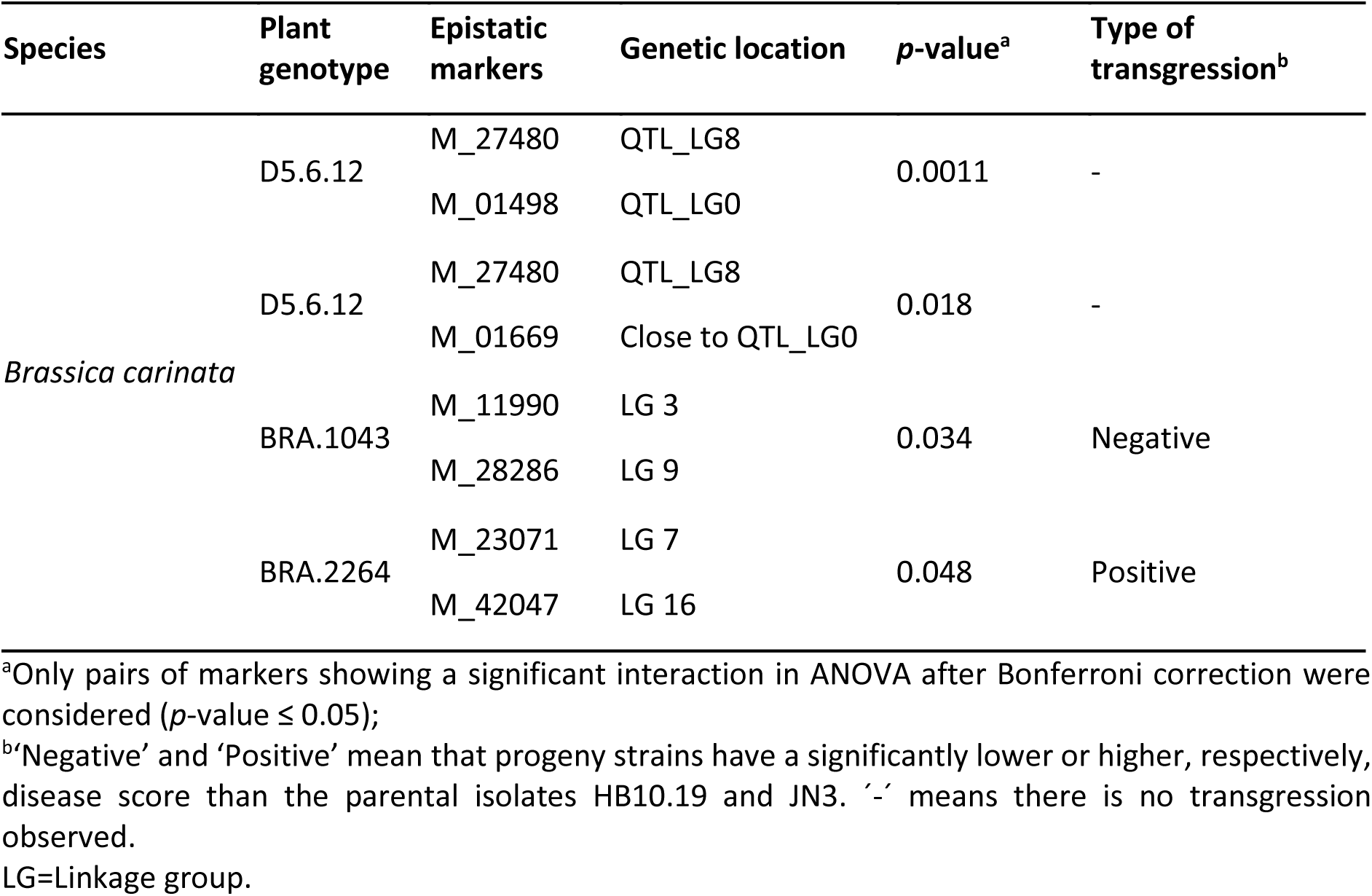
Epistatic relationships between markers influencing *Leptosphaeria maculans* disease score, including transgressive effects, on *Brassica carinata*.

| Species | Plant genotype | Epistatic markers | Genetic location | p-value<sup>a</sup> | Type of transgression<sup>b</sup> |
| --- | --- | --- | --- | --- | --- |
| *Brassica carinata* | D5.6.12 | M_27480 | QTL_LG8 | 0.0011 | - |
|  |  | M_01498 | QTL_LG0 |  |  |
|  | D5.6.12 | M_27480 | QTL_LG8 | 0.018 | - |
|  |  | M_01669 | Close to QTL_LG0 |  |  |
|  | BRA.1043 | M_11990 | LG 3 | 0.034 | Negative |
|  |  | M_28286 | LG 9 |  |  |
|  | BRA.2264 | M_23071 | LG 7 | 0.048 | Positive |
|  |  | M_42047 | LG 16 |  |  |

<sup>a</sup>Only pairs of markers showing a significant interaction in ANOVA after Bonferroni correction were considered ( $p$ -value $\leq 0.05$ );
<sup>b</sup>'Negative' and 'Positive' mean that progeny strains have a significantly lower or higher, respectively, disease score than the parental isolates HB10.19 and JN3. '-' means there is no transgression observed.
LG=Linkage group.

Markers M_27480 and M_01498 exhibited epistasis on *B. carinata* line D5.6.12 (Fig. 4A), and they were the peak markers of the QTL_LG8 and QTL_LG0, respectively. In addition, marker M_27480 showed epistasis with another marker in QTL_LG0, marker M_01669. The interaction between markers M_23071 and M_42047 exhibited positive epistasis, as the aggressiveness of *L. maculans* strains carrying two favourable QTL alleles was higher than expected by a combination of the QTL alleles. The interaction between markers M_11990 and M_28286 is a negative epistasis, as the aggressiveness of *L. maculans* strains carrying two unfavourable QTL alleles was higher than expected a combination of the QTL alleles.

We tested whether the detected epistases could contribute to the transgressions observed in the *L. maculans* progeny. Only allelic combinations i) that were not present in the parental strains, and ii) for which the aggressiveness was lower (or higher) than both parental strains could contribute to the transgression. The four strains exhibiting significant positive transgressive phenotypes on *B. carinata* line BRA.2264 carried an allele combination of the M_23071 and M_42047 epistatic markers that derives from the two parental isolates. These positive transgressive strains carried the favourable allelic combination of these two markers more frequently than the non-transgressive strains (Fisher’s exact test, *p*-value < 0.05; Table 5). This demonstrates that the epistasis between these two markers may, at least in part, explain the positive transgressions observed in the progeny. The two progeny strains exhibiting significant negative transgressive phenotypes on *B. carinata* genotype BRA.1043 carried an allele combination of the M_11990 and M_28286 epistatic markers derived from the two parental isolates. The negative transgressive strains carried the unfavourable allelic combination of these two markers more frequently than the non-transgressive strains (Fisher’s exact test, *p*-value < 0.05; Table 5). This demonstrates that the epistasis between these two markers could be involved, at least in part, in the negative transgressions observed in the progeny.

### Identification of candidate genes in the QTL intervals

We identified 161 and 302 *L. maculans* genes within the QTL intervals detected for pathogenicity on *B. carinata* and *B. napus*, respectively (Table 3; Table S5). We checked their expression profile during the infection of *B. napus* (Gay *et al.,* 2021), the protein polymorphism between the two parental isolates, their location in eu- or heterochromatin during axenic growth (Soyer *et al.,* 2021), and their potential role in pathogenesis (i.e., genes encoding putative effectors or CAZymes; Dutreux *et al.,* 2018; Gay *et al.,* 2021). We considered a gene as a candidate involved in *L. maculans* adaptation toward *B. napus* or *B. carinata* if it encodes a putative or previously described effector or a predicted CAZyme, and additionally exhibits at least one of the following characteristics: i) localization within a TE-rich or heterochromatin region, or ii) expressed by *L. maculans* during early infection of cotyledons of *B. napus* (early, asymptomatic, or switch from biotrophy to necrotrophy stages) as *B. napus* and *B. carinata* are closely related species, and because transcriptomic data describing *L. maculans* expression during infection of *B. carinata* are currently unavailable, or iii) a sequence polymorphism between HB10.19 and JN3. Based on these criteria, we identified candidate genes in all QTL intervals except for QTL_LG9 and QTL_LG18 (Table 6).

**Table 6.**
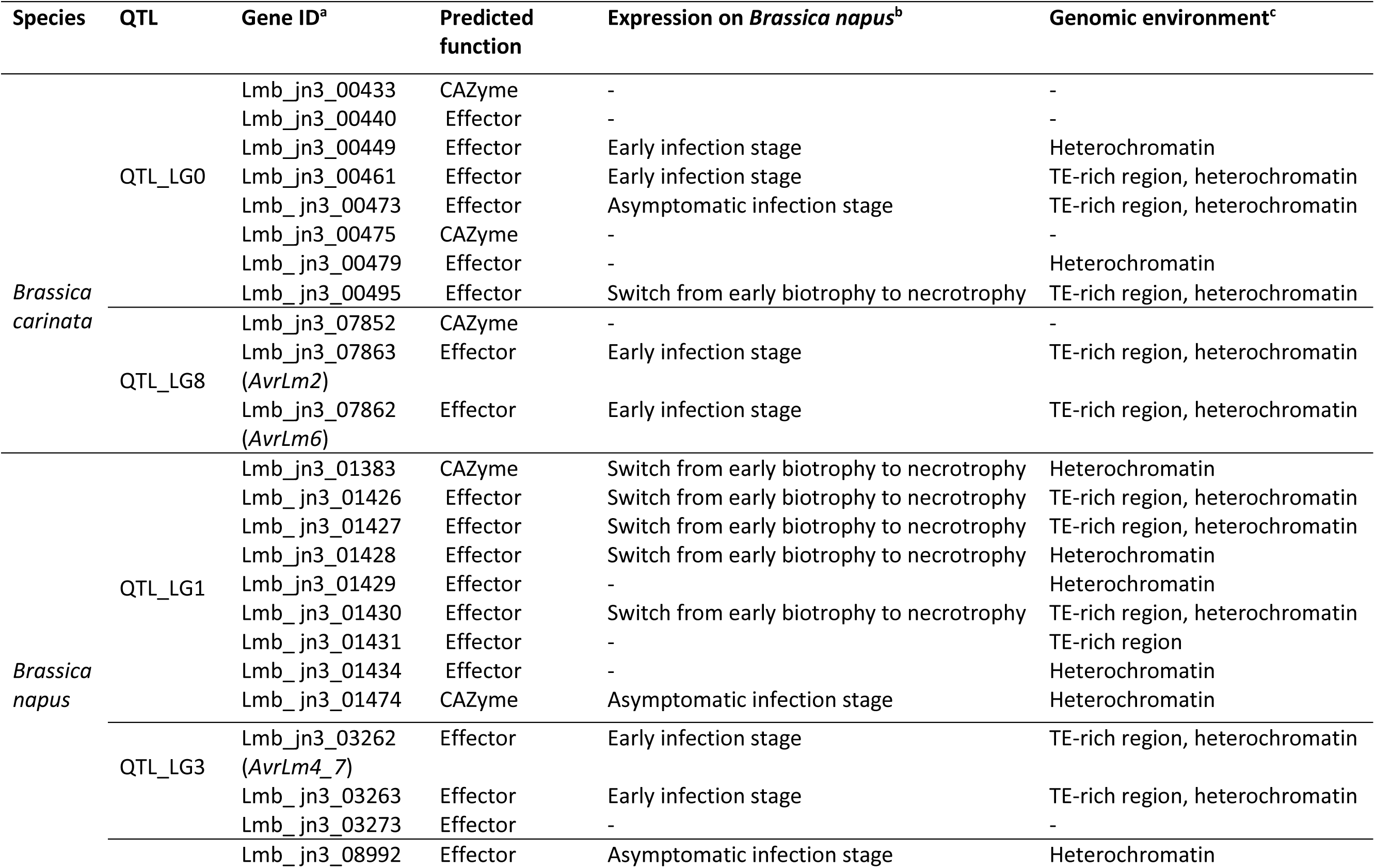
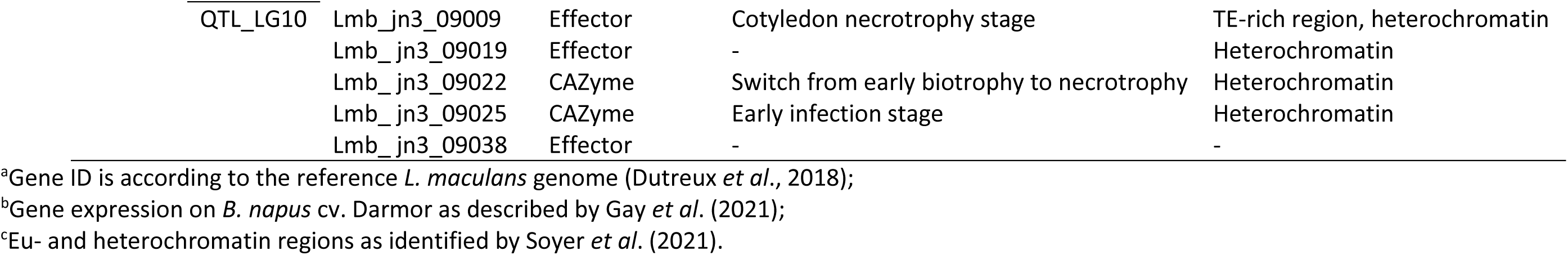
Most relevant candidate genes identified in QTL intervals for pathogenicity of *Leptosphaeria maculans* towards *Brassica napus* and *Brassica carinata*.

We identified 28 genes within the QTL_LG8 confidence interval (the highest phenotypic variance explained for pathogenicity on *B. carinata*), thereby determining pathogenicity in *B. carinata* (Table 3). This region was enriched in genes expressed during *L. maculans* early infection stage of *B. napus* (n=2, *X²* test, *p*-value≤2.6.10^-16^). The QTL_LG8 encompassed two avirulence genes, *AvrLm6* and *AvrLm2*, and the specialized metabolite polyketide synthase 6 (PKS6) located at the peak of the QTL. As the PKS6 gene was not polymorphic between HB0.19 and JN3 (Table S5), it was not selected as a relevant candidate gene. The *AvrLm6* and *AvrLm2* genes are located within TE-rich regions of the *L. maculans* genome. They were highly polymorphic between HB10.19 and JN3 (Table S5), resulting in a distinct protein sequence in HB10.19 and JN3. Moreover, both genes are highly expressed by *L. maculans* during the early infection stage on *B. napus* during a compatible interaction (Table S5). The gene Lmb_jn3_07852 encodes a putative effector and is polymorphic between HB10.19 and JN3. Based on these criteria, *AvrLm2*, *AvrLm6*, and Lmb_jn3_07852 are valuable candidates underlying pathogenicity of *L. maculans* towards *B. carinata* (Table 6).

The confidence interval of QTL_LG0, including 97 genes (Table 3; Table S5), determining pathogenicity towards *B. carinata* was enriched in genes up-regulated by *L. maculans* during the early and asymptomatic stages of cotyledon infection on *B. napus* compared to axenic growth, associated with conidia germination and hyphae penetration of the plant tissue (n=5, *X²* test, *p*-value=≤2.6.10^-16^; Table S5). This interval was also enriched in genes up-regulated by *L. maculans* during the shift from biotrophy to necrotrophy during compatible infection of *B. napus* compared to axenic growth (n=5, *X²* test, *p*-value=≤2.6.10^-16^). Among the seven putative effectors located in QTL_LG0, six of them were selected as candidate genes (Table 6). The putative effector gene Lmb_jn3_00440 was polymorphic between HB10.19 and JN3; gene Lmb_jn3_00449 was up-regulated in *L. maculans* during early infection of *B. napus*; gene Lmb_jn3_00495 was located in TE-rich region and up-regulated during the switch from asymptomatic to necrotrophic infection of cotyledons of *B. napus*; genes Lmb_jn3_00479, Lmb_jn3_00473 and Lmb_jn3_00461 were located in heterochromatin and displayed polymorphism between JN3 and HB10.19; moreover, genes Lmb_jn3_00461 and Lmb_jn3_00473, were up-regulated during early infection of *B. napus*. We also identified the putative CAZyme-encoding genes Lmb_jn3_00433 and Lmb_jn3_00475, which showed nucleotide polymorphisms between HB10.19 and JN3 (Table S5). Based on these criteria, eight genes are valuable candidates underlying pathogenicity of *L. maculans* towards *B. carinata* (Table 6).

The confidence interval of QTL_LG3 (responsible for the highest phenotypic variance for the pathogenicity on *B. napus*) contained 27 genes. It was located in a subtelomeric region and was enriched in genes up-regulated during the *L. maculans* early and asymptomatic stages of infection in *B. napus* compared to axenic growth (n=3, *X²* test, *p*-value=≤2.6.10^-16^). Among the four effector genes in this QTL interval, we identified the avirulence gene *AvrLm4-7*, located within a TE-rich, heterochromatic region in JN3, absent in isolate HB10.19 compared to JN3, and expressed by *L. maculans* during early infection stage on *B. napus* (Table S5; Table 6). The gene Lmb_jn3_03263 encoding a putative effector was also considered a good candidate as it was located in a TE-rich region, and up-regulated during a compatible infection of *B. napus* (Table 6). Finally, the putative effector Lmb_jn3_03273 exhibited polymorphism between HB10.19 and JN3. Based on these criteria, we identified three genes as valuable candidates underlying pathogenicity of *L. maculans* towards *B. napus* (Table 6).

QTL_LG1 detected on *B. napus*, encompassing 135 genes (Table 3), is enriched in genes up-regulated by *L. maculans* during the asymptomatic and switch from biotrophy to necrotrophy stages of *B. napus* infection compared to axenic growth (n=8, *X²* test, *p*-value<2.2x10^-16^; Table S5). Among the 13 putative effector encoding genes located in this interval, four are associated with heterochromatin domains and up-regulated during the shift from asymptomatic to necrotrophic infection of cotyledons of *B. napus* (genes Lmb_jn3_01426, Lmb_jn3_01427, Lmb_jn3_01428, and Lmb_jn3_01430) with Lmb_jn3_01430 exhibiting polymorphism between HB10.19 and JN3 (Table S5). Putative effector gene Lmb_jn3_01431 exhibited sequence polymorphism between HB10.19 and JN3 and was located in a TE-rich region; genes Lmb_jn3_01434 and Lmb_jn3_01429 were located in heterochromatin region (Table S5). Also, QTL_LG1 encompassed three putative CAZyme encoding genes, among which gene Lmb_jn3_01383 was located in heterochromatin and up-regulated by *L. maculans* during the switch between the asymptomatic and the necrotrophic stage of cotyledon infection, and Lmb_jn3_01474 was up-regulated by *L maculans* during the asymptomatic stage of infection and exhibited polymorphism between HB10.19 and JN3. Based on these criteria, we considered these nine genes as good candidates to explain the pathogenicity of *L. maculans* towards *B. napus* (Table 6).

The interval of QTL_LG10 detected during infection on *B. napus*, encompassing 76 genes (Table 3), and located in a subtelomeric region, was enriched in genes associated with heterochromatin-domain (n=21, *X²*, *p*-value<2.2x10^-16^). This interval included four putative effector genes. Among them, three are located in heterochromatin: genes Lmb_jn3_08992, Lmb_jn3_09019, and Lmb_jn3_09009, with gene Lmb_jn3_08992 up-regulated during the asymptomatic stage of infection of *B. napus* and polymorphic between HB10.19 and JN3; and gene Lmb_jn3_09009, located in a TE-rich region and exhibiting polymorphism between HB10.19 and JN3. The fourth putative effector gene, Lmb_jn3_09038, exhibited polymorphism between HB10.19 and JN3. Two genes encoded putative CAZymes, *i.e.*, genes Lmb_jn3_09022 and Lmb_jn3_09025 were located in heterochromatin region, and up-regulated during early infection of *B. napus*, with the former exhibiting polymorphism between HB10.19 and JN3 (Table S5). Based on these criteria, we considered these six genes as good candidates to explain the pathogenicity of *L. maculans* towards *B. napus* (Table 6).

Overall, we identified 11 and 18 candidate genes for the pathogenicity of *L. maculans* towards *B. carinata* and *B. napus*, respectively.

## Discussion

In this study, we aimed to identify the genomic regions of *L. maculans* involved in the adaptation towards *B. napus* and/or *B. carinata*. The identification of determinants of fungal host spectra remains elusive, and the factors involved in fungal adaptation are of major interest for preventing fungal bypass of host resistance. We generated an ultra-dense linkage map of *L. maculans* covering 92.1% of the *L. maculans* reference genome. This genetic map consists of 20 linkage groups (corresponding to the number of supercontigs of *L. maculans*), comprising a total of 34,147 SNP markers distributed in 1,466 unique positions (also called ‘bins’ represented by skeleton markers). Previously, Dutreux *et al*. (2018) used a cross between two geographically distant *L. maculans* isolates adapted to *B. napus* to construct a genetic map of 20 linkage groups, comprising a total of 20,103 SNPs. Although both genetic linkage maps have 20 linkage groups corresponding to the number of chromosomes of *L. maculans* (Dutreux *et al*., 2018), we identified a higher number of SNP between the two parental strains (JN3 and HB10.19), which indicates a higher genetic distance between these strains compared to JN2 and NzT4, the parents of the cross used in Dutreux *et al*. (2018). Few genetic maps of *L. maculans* are presently available. Pongam *et al*. (1998) were the first to build a genetic map of *L. maculans* using 56 AFLP (Amplified Fragment Length Polymorphism) markers. Nine linkage groups covering a total genome length of 340 cM were detected. Cozjinsen *et al*. (2000) generated a genetic map comprising 155 AFLP markers and three RAPD (Random Amplified Polymorphic DNA) markers. This genetic map contained 21 linkage groups and covered 1,520 cM. Performing three *L. maculans* crosses and combining AFLP, RAPD, ISSR (Inter Simple Sequence Repeat), and Rep-PCR (Repetitive Polymerase Chain Reaction) markers, Kuhn *et al*. (2006) obtained three genetic maps encompassing 78, 35, and 61 linkage groups, with total sizes of 2,219, 430, and 867 cM, respectively. Ghanbarnia *et al*. (2012) generated a genetic linkage map to identify the locus of the avirulence gene *AvrLepR1*. They used 283 informative sequence-related amplified polymorphism (SRAP) markers to generate a genetic map of 2,076.4 cM, comprising 36 linkage groups. These previous studies used a limited number of markers, identified a large number of linkage groups, and based their analysis on the outdated estimation of *L. maculans* genome size (34 Mb in Cozijnsen *et al.,* 2000). In comparison with these older studies, the greater coverage and length of our genetic map, or that of Dutreux *et al*. (2018) is due to a dense marker coverage resulting from whole genome sequencing of progeny, and a great number of progeny strains (except for Kuhn *et al*. (2006), where 98, 42, and 46 progeny strains from three different crosses were combined to build the genetic map) obtained for the cross between JN3 and HB10.19. This emphasises the usefulness of whole-genome sequencing for generating ultra-dense genetic maps.

In eukaryotes, there are fewer crossovers in heterochromatin and TE-rich regions (Mézard *et al*., 2015; Kent *et al*., 2017; Hartmann *et al*., 2021b), and recombination hotspots are observed in GC-rich regions (Croll *et al*., 2015; Sonnenberg *et al*., 2017; Laurent *et al*., 2018). The reference genome of *L. maculans* harbours one TE-rich dispensable chromosome (Leclair *et al.,* 1994; Balesdent *et al*., 2013), which is present in the parental isolates of this study, JN3 and HB10.19. The dispensable chromosomes follow a non-Mendelian inheritance (Komluski *et al*., 2022). In *L. maculans*, it was previously estimated that 3.5 to 6.7% of the progeny lack the dispensable chromosome after meiosis (Balesdent *et al.,* 2013). In our study, we found 12.6% of the progeny strains lacking the dispensable chromosome. We found that the dispensable chromosome had a few crossovers, with only six progeny strains showing recombination on it. In *L. maculans*, 92.5% of the dispensable chromosome is composed of TE compared to 33% for the core chromosomes (Rouxel *et al.,* 2011). We corroborated previous results identifying non-Mendelian segregation of the dispensable chromosome of *L. maculans* (Leclair *et al.,* 1994; Balesdent *et al.,* 2013) and fewer crossovers in TE-rich regions.

We observed a continuum of aggressiveness in the progeny strains on the two *Brassica* species. This suggests that the adaptation of *L. maculans* towards both *Brassica* species is under polygenic control. We observed 11 and 12 progeny strains causing symptoms on two genotypes of *B. carinata* (D.5.612 and BRA.1043), corresponding to an approximate avirulence/virulence ratio of 8:1, as *L. maculans* is a haploid species. This ratio could be explained by the segregation of three unlinked genes in the progeny, with only strains carrying the three recessive alleles being virulent. In accordance with this result, the linkage map analysis enabled us to detect three specific QTL intervals for the pathogenicity of *L. maculans* towards *B. carinata*, while four QTL intervals were detected for pathogenicity towards *B. napus*. The three QTL identified for *L. maculans* adaptation to *B. carinata* showed an overall additive effect, and these QTL explained a high proportion of the variation in the disease score of the progeny. These results support the hypothesis of a polygenic adaptation of *L. maculans*, facilitating its infection of *B. carinata*. Altogether, these results suggest that a relatively small number of genes are involved in *L. maculans* adaptation to *B. carinata*.

None of the intervals identified is involved in adaptation to both *Brassica* species. Both species-specific and cultivar-specific QTL have been identified. The QTL_LG8 was detected for the pathogenicity of *L. maculans* towards the five *B. carinata* genotypes used in the study. Similarly, QTL_LG3, which determines pathogenicity of *L. maculans* on *B. napus*, was detected across all three genotypes of *B. napus*. This suggests that these QTL_LG8 and QTL_LG3 are species-specific and are robust genomic regions involved in the adaptation of *L. maculans* to any genotype of *B. carinata* or *B. napus*, respectively. All the other QTLs identified were detected for only certain genotypes, suggesting they are genotype-specific.

QTL_LG8 and QTL_LG3 explained the majority of the observed phenotypic variation for the pathogenicity towards *B. carinata* and *B. napus*. They are enriched in TE-rich regions that cover 83 and 63% of the genomic region, respectively, and both correspond to subtelomeric regions. Our identification of the genomic regions involved in the adaptation of *L. maculans* toward its host, *B. napus*, corroborated previous findings indicating that plastic regions (and especially subtelomeric regions) are frequently involved in host adaptation of fungi (Rouxel *et al*., 2011; Grandaubert *et al*., 2014; Faino *et al*., 2016; Seidl and Thomma, 2017). Our analysis highlighted that, as for adaptation of *L. maculans* to *B. napus* hosts, pathogenicity toward *B. carinata*, a nonhost species of *L. maculans*, also involves genes embedded in TE-rich regions. This is also found in the genomic regions involved in adaptation to different host species in fungi, such as *Ceratocystis* species. *Ceratocystis fimbriata* is a fungus that infects only sweet potato (Valdetaro *et al*., 2019), while *Ceratocystis manginecans* infects various tree species, such as the acacia tree (Tarigan *et al*., 2011), yet is not adapted to sweet potato (Baker *et al*., 2003; Fourie *et al*., 2018). An interspecific cross was performed between these two *Ceratocystis* species, and a QTL analysis of aggressiveness on both sweet potato and acacia was realized (Fourie *et al.,* 2019). Two QTL regions underlying aggressiveness on sweet potato were located in TE-rich regions. Our results suggest that plastic regions are likely involved in the adaptation of *L. maculans* to both *Brassica* host and nonhost species.

We observed phenotypic transgressions in the progeny of *L. maculans*. Such transgressive phenotypes are rarely reported in fungi, whereas they have been well documented in plants (Rieseberg *et al*., 1999, 2003; Stelkens and Seehausen, 2009). However, few studies reported positive or negative phenotypic transgressions in fungi (Chung *et al*., 1997; Cumagun *et al*., 2004; Voss *et al*., 2010; Gibson *et al*., 2014; Gohari *et al*., 2015; Caffier *et al*., 2022). The transgressions observed in our progeny, i.e., generation of fungal genotypes adapted to both *Brassica* species, support the hypothesis that sexual reproduction, with recombination of loci during meiosis, favours fungal adaptation or host jumps (McDonald and Linde, 2002; Thines, 2019). We identified epistasis between two marker pairs as part of the explanation for the transgression. The simplest genetic model would be a combination of favourable (or unfavourable, depending on the direction of the transgression) alleles originating from the two parental isolates (“additive” model). We rejected this model because all favourable alleles at the detected QTL were present in the best-adapted parental isolate, whatever the host genotype. The other classical genetic model to explain transgressions is the “epistasis” model, in which a combination of alleles originating from the two parental isolates confers a more-than-additive effect (or less-than-additive depending on the transgression direction). Accordingly, we identified epistasis between two non-additive marker pairs as being part of the transgression explanation. This study is the first to describe epistatic markers involved in phenotypic transgression and adaptation of a fungus to a nonhost plant. This appears to be genotype-specific, as epistasis between markers was significant only on some *B. carinata* genotypes. Unfortunately, we were unable to explain the transgression with epistasis between markers for transgressive strains for interaction with all the other *B. carinata* and *B. napus* genotypes. This issue could be due to insufficient statistical power to detect epistasis in our analyses or because the epistases involved were more complex (i.e., involving more than two markers). The fact that epistasis between non-additive markers is a major genetic component of phenotypic transgressions and host adaptation in a fungal progeny makes it very difficult to predict in which conditions such transgressions can be observed. Indeed, these epistases are strictly dependent on the two parental isolates crossed to generate the progeny, and they cannot be predicted from the identified additive QTL.

QTL intervals involved in the interaction of *L. maculans* with *B. carinata* and *B. napus* contained 162 and 303 genes, respectively. Effector genes, up-regulated during host infection, are involved in host adaptation and pathogenesis (Schulze-Lefert and Panstruga, 2011; Lo Presti *et al*., 2015; Sánchez-Vallet *et al*., 2018; Gay *et al*., 2021). Similarly, genes up-regulated during *B. napus* infection compared to *in vitro* conditions are supposed to be involved in pathogenesis toward the plant host, such as CAZymes, which are up-regulated by *L. maculans* at specific stages of infection (Gay *et al*., 2021; 2023). In addition, genes located in plastic regions are more prone to deletions in *L. maculans* (Rouxel and Balesdent, 2017), which allows the fungus to escape recognition by the immune system of *B. napus*. Hence, to identify the best candidate genes involved in adaptation towards both *Brassica* species, we focused on effectors and CAZymes, their polymorphism between the two parental isolates JN3 and HB10.19, their expression during infection on *B. napus* (Gay *et al.,* 2021), and their location in plastic genomic regions (heterochromatic and TE-rich regions; Soyer *et al*., 2021). In total, we identified 18 candidate genes that may be involved in the adaptation of *L. maculans* to the two *Brassica* species. The QTL_LG8, specific to pathogenicity towards *B. carinata*, explains up to 67.6% of the phenotypic variation for the disease score, suggesting that a single gene, possibly involved in a gene-for-gene relationship, could be sufficient to explain this phenotypic trait. We identified the effectors within this QTL_LG8 as good candidates for the adaptation of *L. maculans* to *B. carinata*. Functional analysis will help decipher their implications in the observed adaptation.

Altogether, our results suggest a complex adaptation of *L. maculans* to different *Brassica* species, involving additive QTL, epistasis between non-additive markers, plastic genomic regions, and possibly gene-for-gene relationships.

## CRediT authorship contribution statement

**Julie M. Noah**: Investigation, Methodology, Formal analysis, Validation, Visualization, Writing – original draft, Writing – review and editing; **Marie-Hélène Balesdent**: Conceptualization, Funding acquisition, Validation, Writing – review and editing; **Marie Foulongne-Oriol**: Writing – review and editing; **Mathilde Gorse**: Formal analysis, Investigation, Writing – review and editing; **Camilla Langlands-Perry**: Methodology, Writing – review and editing; **Nicolas Lapalu**: Data curation, Methodology, Writing – review and editing; **Thierry C. Marcel**: Methodology, Writing – review and editing, **Benoit Moury**: Methodology, Writing – review and editing, **Thierry Rouxel**: Conceptualization, Funding acquisition, Validation, Writing – review and editing, **Jessica L. Soyer**: Conceptualization, Funding acquisition, Project administration, Supervision, Validation, Writing – review and editing

## Data availability

The data that support the findings of this study are openly available under SRA BioProject ID PRJEB39398 and PRJNA1128709.

## Acknowledgements

This work was funded by the French National Research Agency project Gandalf (ANR-12-ADAP-0009) and a grant from the INRAE department “Santé des Plantes et Environnement (SPE)” (Project “Phytoadapt”). Julie M. Noah benefits from a grant from the INRAE SPE department and the “Club Phoma” consortium. The authors particularly thank Laurent Coudard (BIOGER) for the management of *L. maculans* isolates, the greenhouse workers for plant management, the administrative supporting staff for the administrative and financial follow-up of this project, the laboratory glassware management staff of the BIOGER research unit, and members of the “Effectors and Pathogenesis of *L. maculans*” group for fruitful discussions. The EPLM group benefits from the support of Saclay Plant Sciences-SPS (ANR-17-EUR-0007).

## Supplementary tables

**Table S1. Summary of the crossovers detected in the progeny issued from the cross between JN3 and HB0.19 isolates of *Leptosphaeria maculans***

This analysis was realised using the 92 progeny strain sequences used to generate the genetic linkage map.

^a^Number of crossovers for each linkage group among the strains of the progeny;

^b^Crossover density was calculated by dividing the crossover number by the physical size of each LG; ^c^For LG 19, we calculated the mean crossover count as the total number of crossovers detected in the progeny divided by the number of progeny strains carrying the dispensable chromosome (n = 80).

**Table S2. Percentage of virulence of the HB10.19 and JN3 parental isolates and the V77 progeny strains on *Brassica napus* and *Brassica carinata* cotyledons**

**Table S3. Details of the QTL identified for each of the *Brassica napus* genotypes and the different phenotypic traits assessed**

ᵃGenotype of *B. napus* on which the phenotypes were measured;

ᵇBiological replicates performed at different times. The replicates and phenotypes that did not allow to identify a QTL are not shown in this table;

^c^Indicates the parental allele (JN3 or HB10.19) which confers higher pathogenicity; QTL_LGx, indicates QTL identified on linkage group (LG) x.

**Table S4. Details of the QTL identified for each of the *Brassica carinata* genotypes and the different phenotypic traits assessed**

ᵃGenotype of *B. carinata* on which the phenotypes were measured;

ᵇBiological replicates performed at different times. The replicates and phenotypes that did not allow to identify a QTL are not shown in this table;

^c^Indicates the parental allele (JN3 or HB10.19) which confers higher pathogenicity; QTL_LGx, indicates QTL identified on linkage group (LG) x.

**Table S5. Genes included in the QTL intervals involved in the adaptation of Leptosphaeria maculans to *Brassica napus* and *Brassica carinata***

^a^Genome from Dutreux *et al*. (2018);

^b^GO annotation of *L. maculans* genes were retrieved from Durtreux *et al*. (2018);

^c^Genes encoding putative CAZyme, proteinaceous effector, PKS or NRPS (Gay *et al.,* 2021);

^d^Genes associated with H3K4me2, H3K9me3 or H3K27me3 during axenic culture in the wild type strain (Soyer *et al*., 2021);

^e^Description of the stage at which genes of *L. maculans* are up-regulated during infection of *B. napus* (Gay *et al*., 2021);

^f^Mutations identified in HB10.19 relative to JN3 are indicated; only non-synonymous mutations are specified

## Supplementary figures

**Fig. S1.**
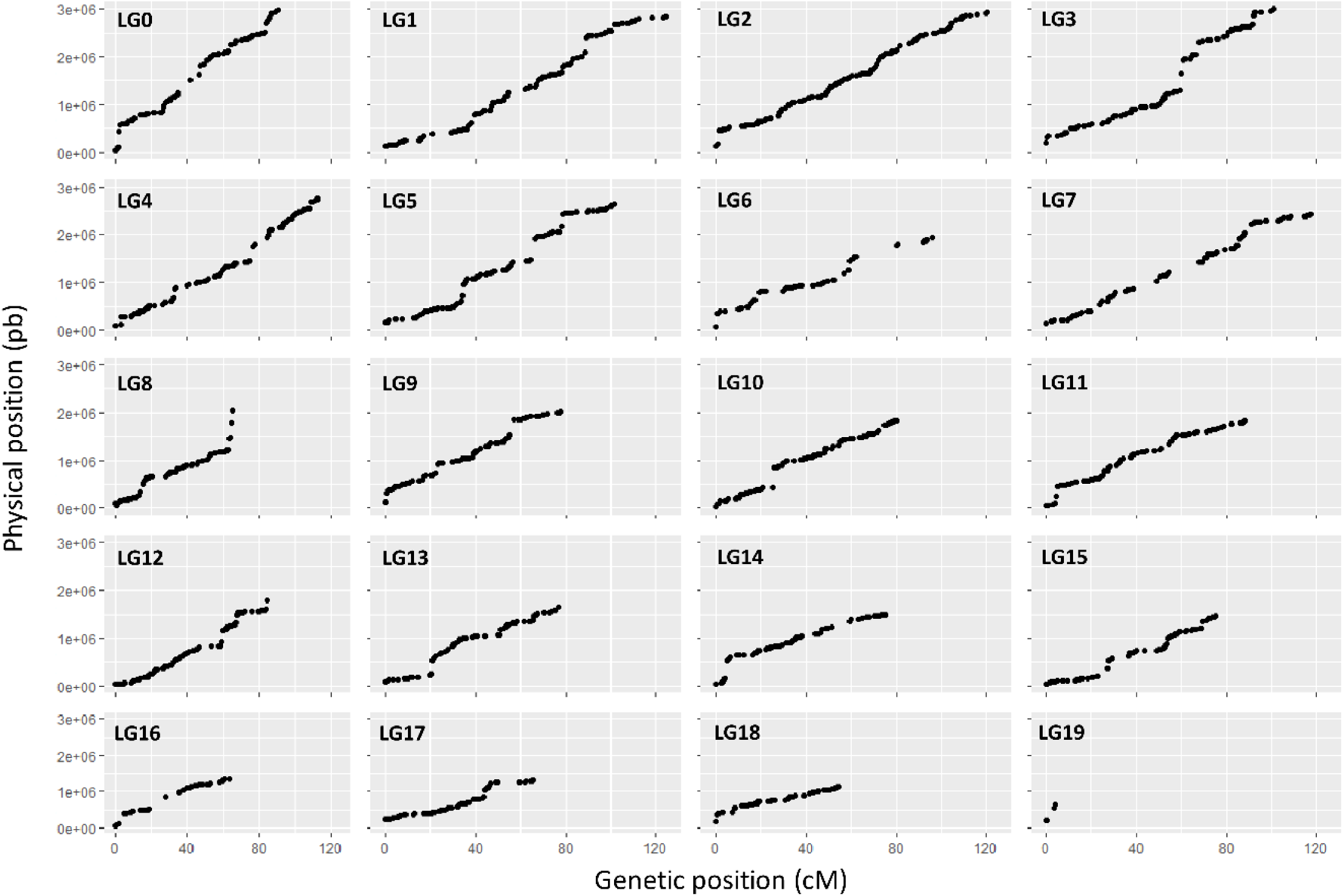
Relationship between the genetic markers (cM) and their physical position (pb) along the *Leptosphaeria maculans* genome. Each linkage group (LG) corresponds to a chromosome of the *L. maculans* genome and LG19 corresponds to the dispensable chromosome.

**Fig. S2.**
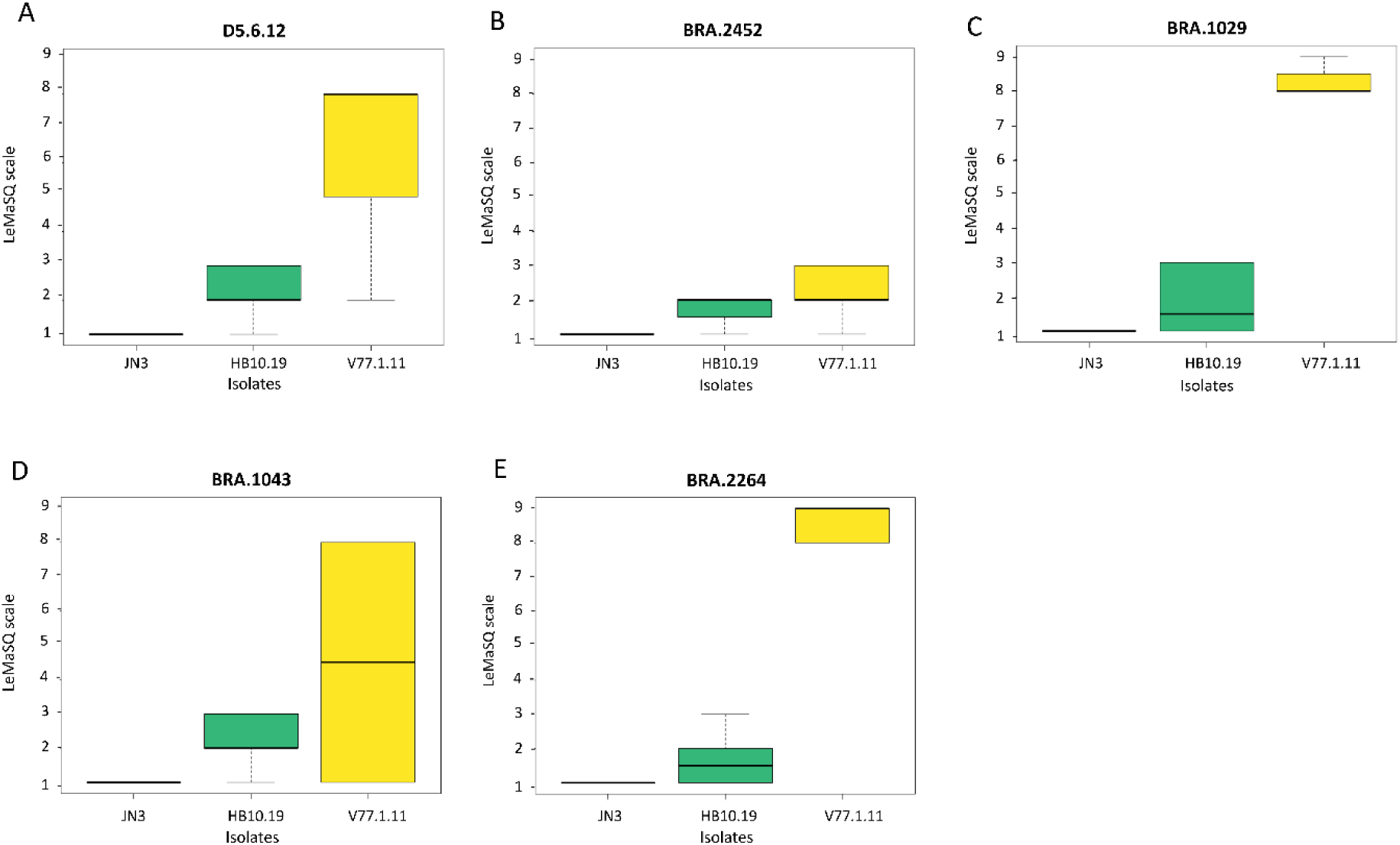
Assessment of symptoms induced by *Leptosphaeria maculans* isolates JN3 and HB10.19, and the V77.1.11 progeny strain on cotyledons of five *Brassica carinata* lines. Symptoms on cotyledons were evaluated following the LeMaSQ scale.

**Fig. S3.**
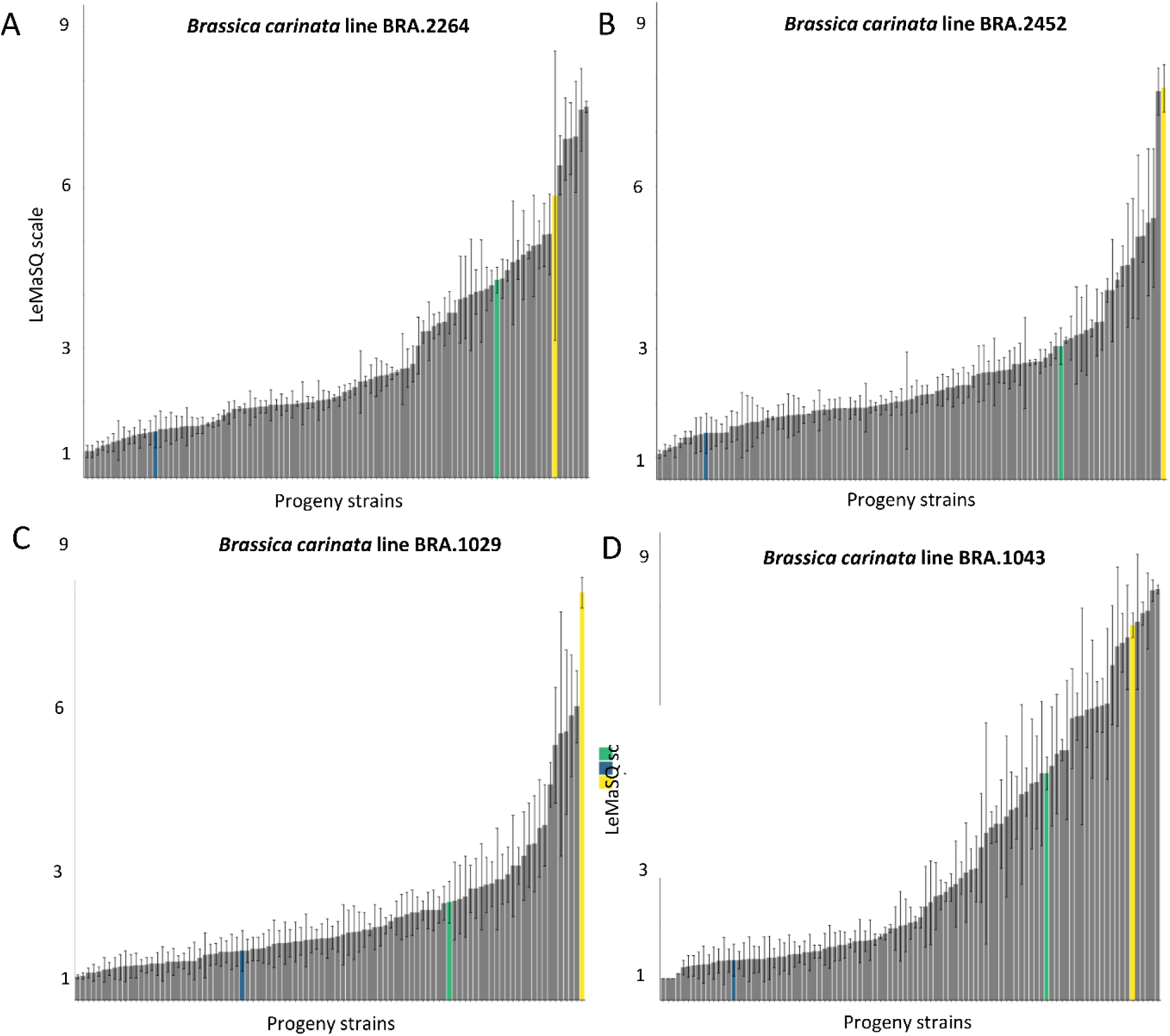
Distribution of disease scores caused by the *Leptosphaeria maculans* V77 progeny strains on cotyledons of four *Brassica carinata* lines. Symptoms were evaluated following the LeMaSQ scale. Dark blue bar, isolate JN3; green bar, isolate HB10.19; grey bars, V77 progeny strains, with the V77.1.11 strain highlighted in yellow. Thin bars represent the standard deviation for each strain.

**Fig. S4.**
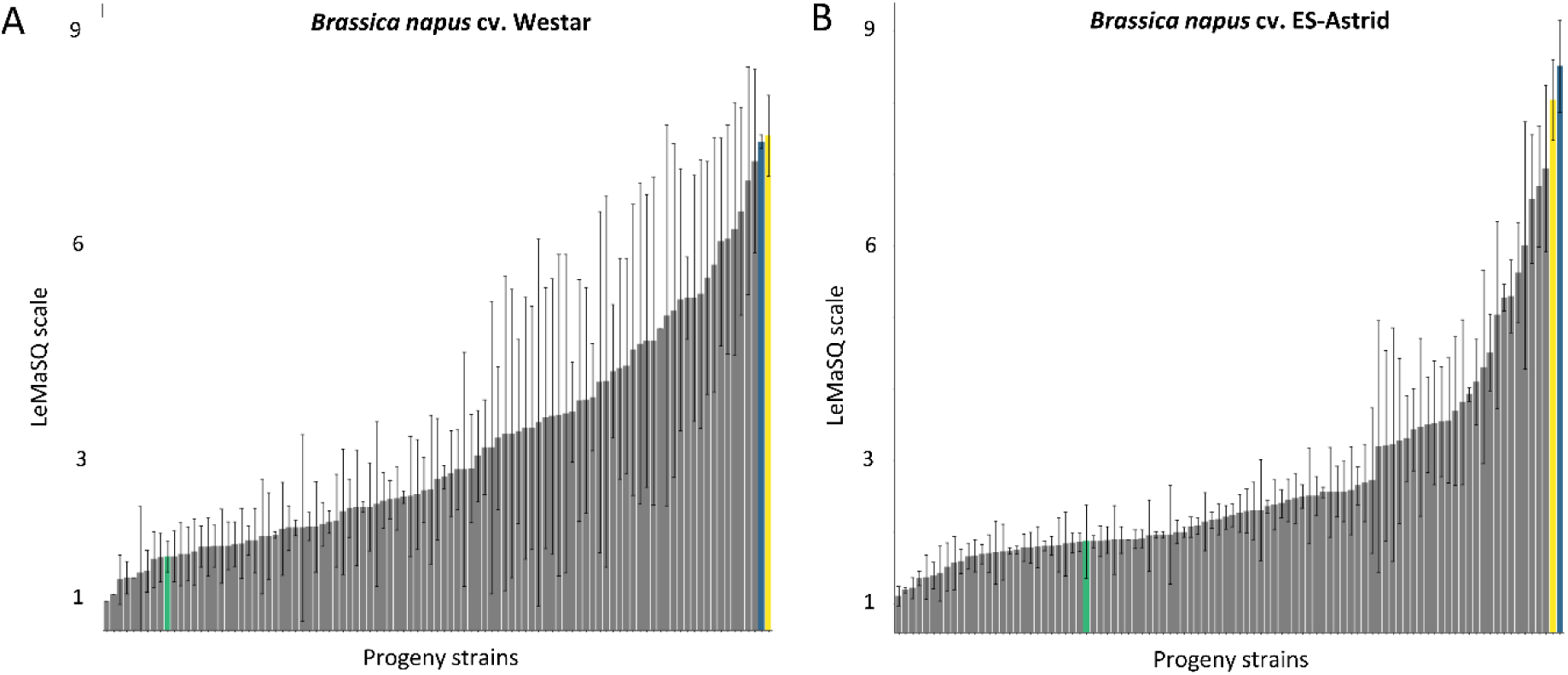
Distribution of disease scores caused by the *Leptosphaeria maculans* V77 progeny strains on cotyledons of two *Brassica napus* cultivars. Symptoms were evaluated following the LeMaSQ scale. Dark blue bar, isolate JN3; green bar, isolate HB10.19; grey bars, V77 progeny strains, with the V77.1.11 strain highlighted in yellow. Thin bars represent the standard deviation for each strain.

**Fig. S5.**
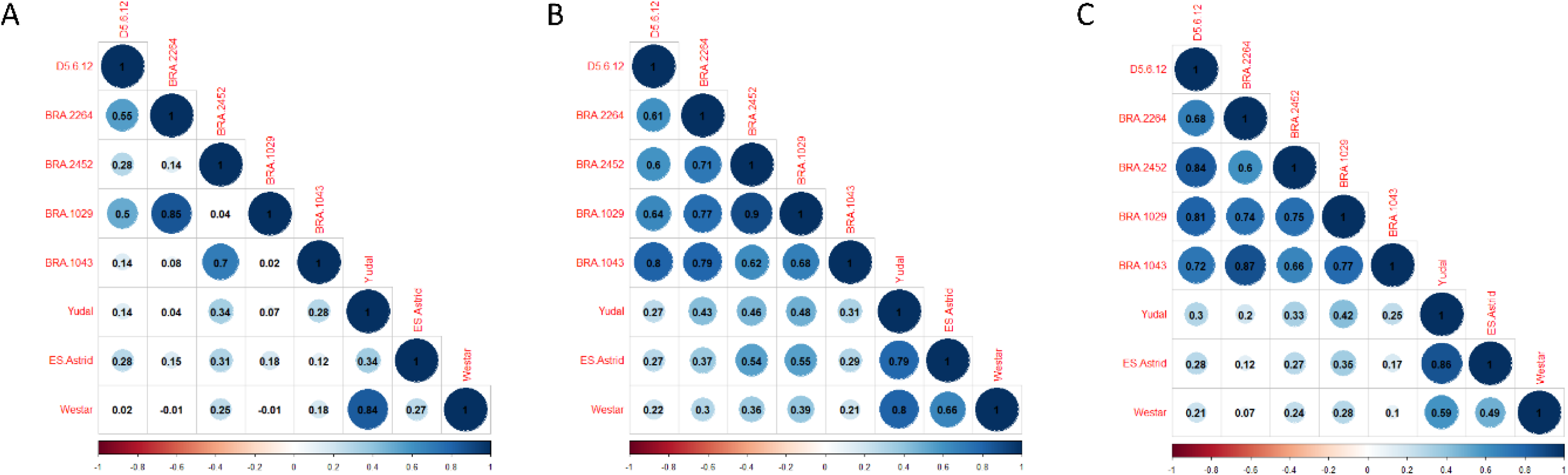
Pearson’s coefficients of correlation between the *Brassica* spp. genotypes for the phenotypes assessed, i.e., the A) disease score, B) virulence percentage, and C) symptom area induced by strains of the V77 progeny. Blue colours represent a positive correlation, and red colours represent a negative correlation. The correlation coefficient is indicated in each circle. D5.6.12, BRA.2264, BRA.2452, BRA.1029, and BRA.1043 are *Brassica carinata* genotypes, and Yudal, ES-Astrid, and Westar are *Brassica napus* cultivars. Non-significant correlations are represented on a white background (no circle).

**Fig. S6.**
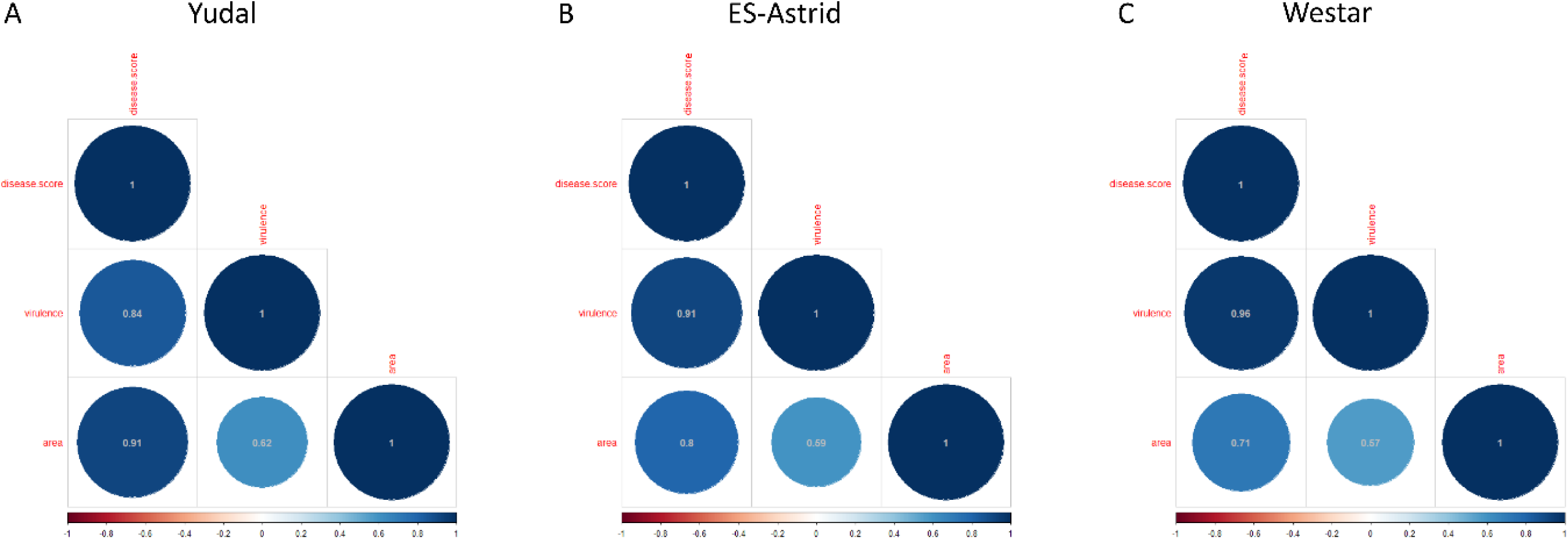
Pearson’s coefficients of correlation between the phenotypes assessed, i.e., the disease score, virulence percentage, and symptom area induced by strains of the V77 progeny on *Brassica napus* cvs. A) Yudal, B) ES-Astrid, and C) Westar. Blue colours represent a positive correlation, and red colours represent a negative correlation. The correlation coefficient is indicated in each circle. disease.score: disease score evaluated according to the LeMaSQ scale, virulence: virulence percentage, area: symptom area.

**Fig. S7.**
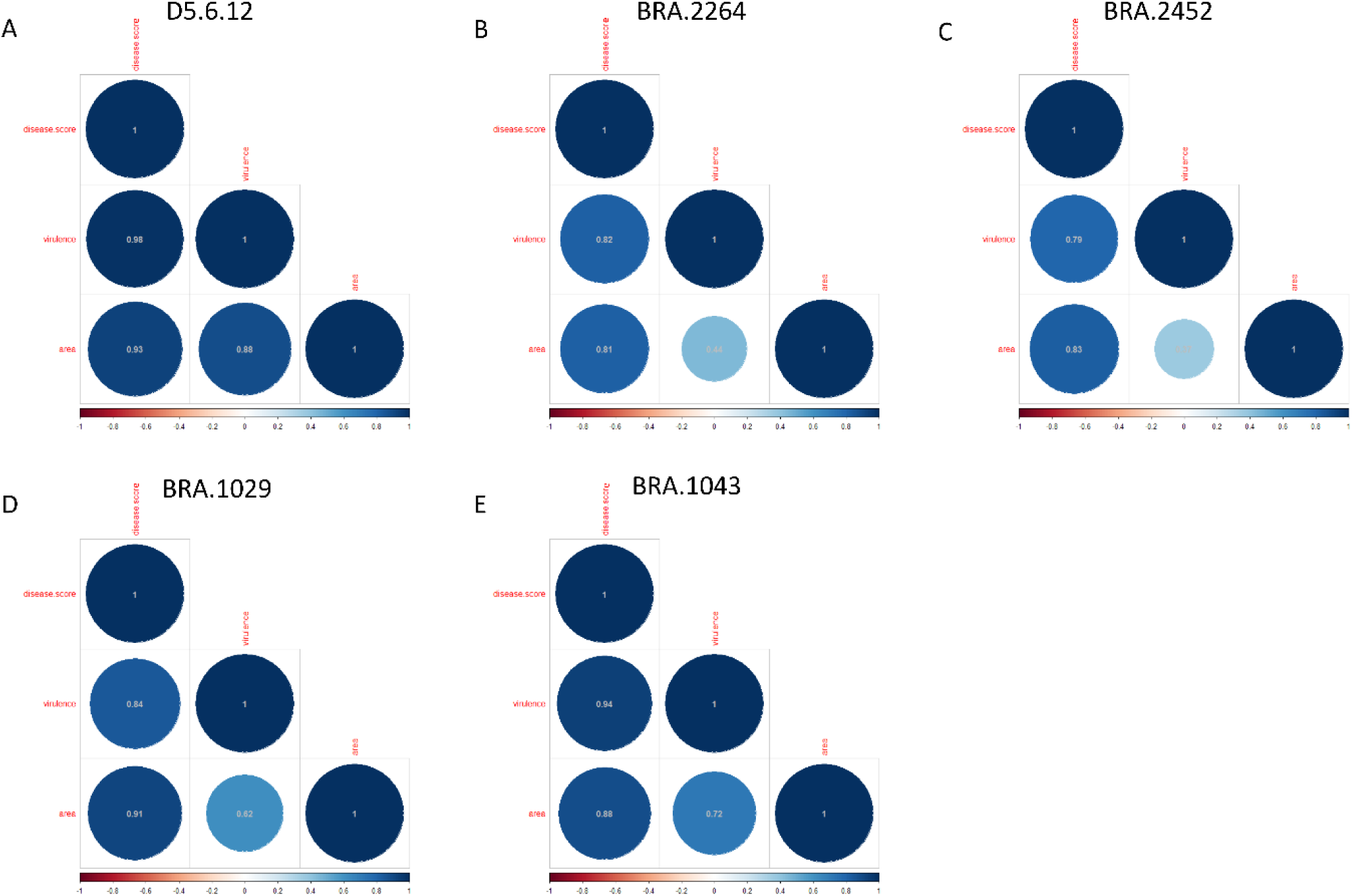
Pearson’s coefficients of correlation between the phenotypes assessed, i.e, the disease score, virulence percentage, and symptom area induced by the V77 progeny on *Brassica carinata* lines A) D5.6.12, B) BRA.2264, C) BRA.2452, D) BRA.1029 and E) BRA.1043. Blue colours represent a positive correlation, and red colours represent a negative correlation. The correlation coefficient is indicated in each circle. disease.score: disease score evaluated according to LeMaSQ scale, virulence: virulence percentage, area: symptom area.

**Fig. S8.**
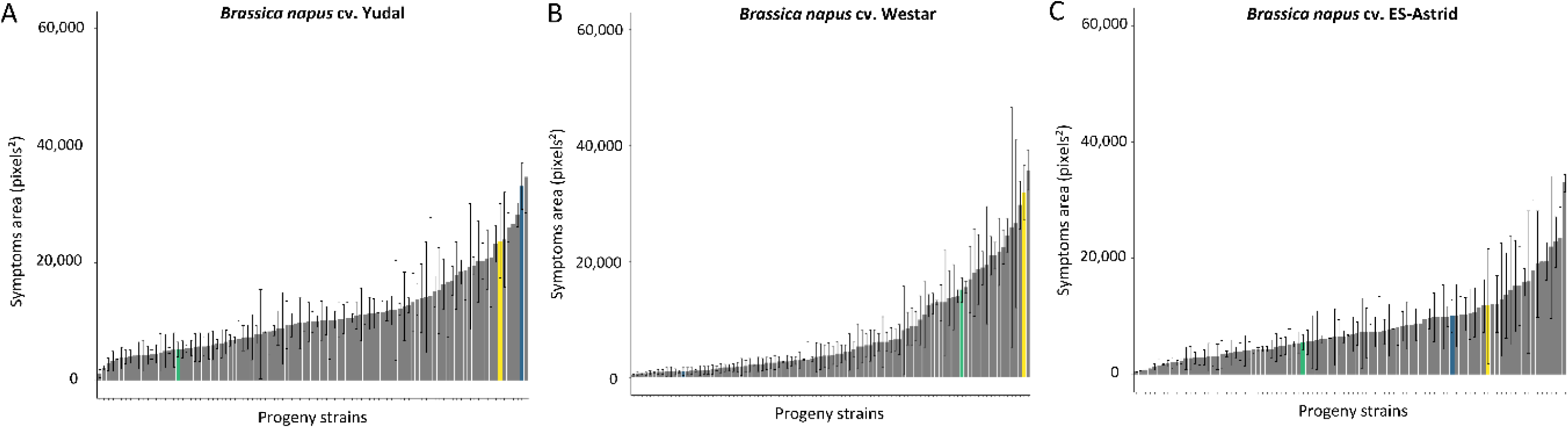
Distribution of symptom area caused by the *Leptosphaeria maculans* V77 progeny strains on *Brassica napus* cotyledons. Dark blue bar, isolate JN3; green bar, isolate HB10.19; grey bars, V77 progeny strains, with the V77.1.11 strain highlighted in yellow. Thin bars represent the standard deviation for each strain.

**Fig. S9.**
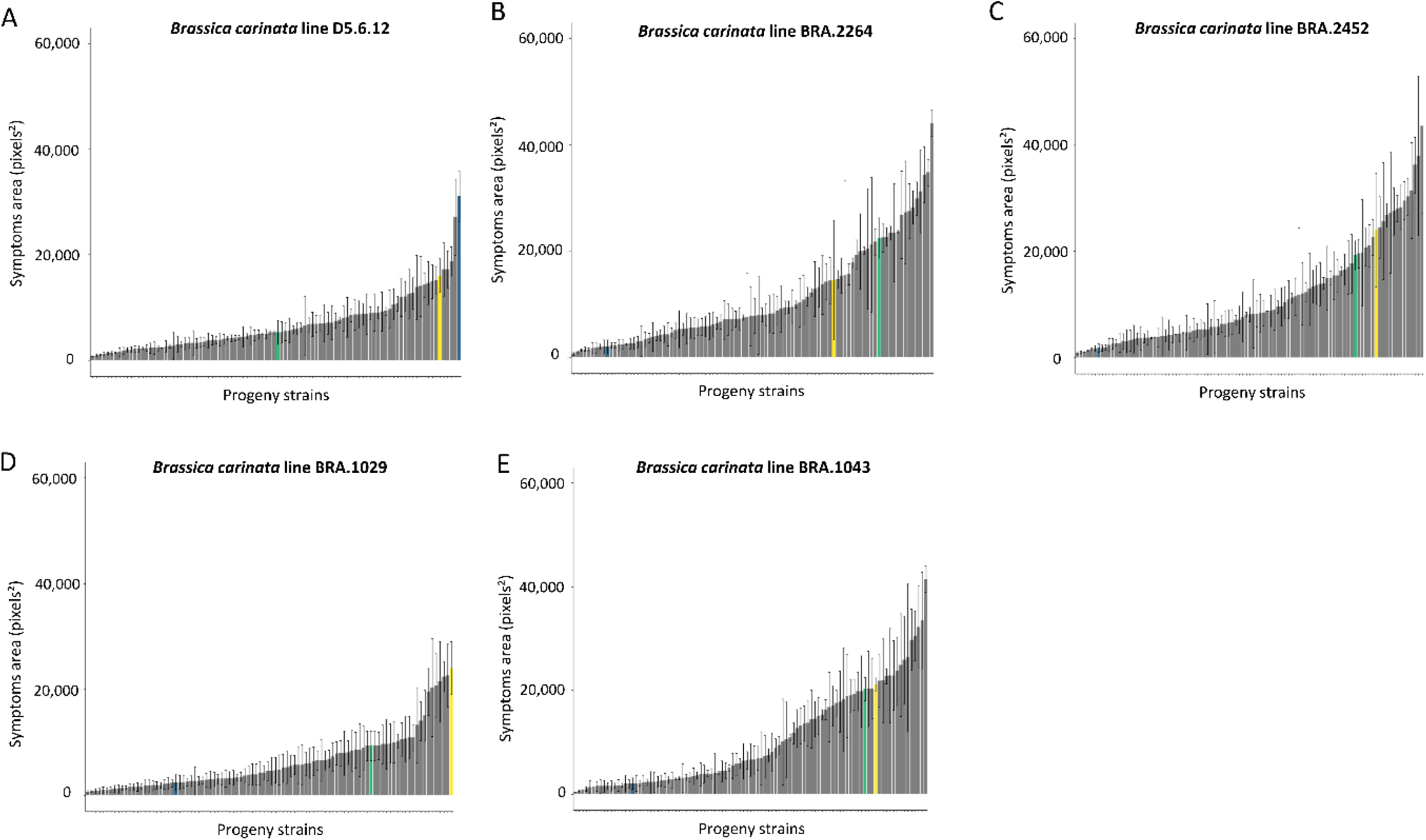
Distribution of symptom area caused by the *Leptosphaeria maculans* V77 progeny strains on *Brassica carinata* cotyledons. Dark blue bar, isolate JN3; green bar, isolate HB10.19; grey bars, V77 progeny strains, with the V77.1.11 strain highlighted in yellow. Thin bars represent the standard deviation for each strain.

**Fig. S10.**
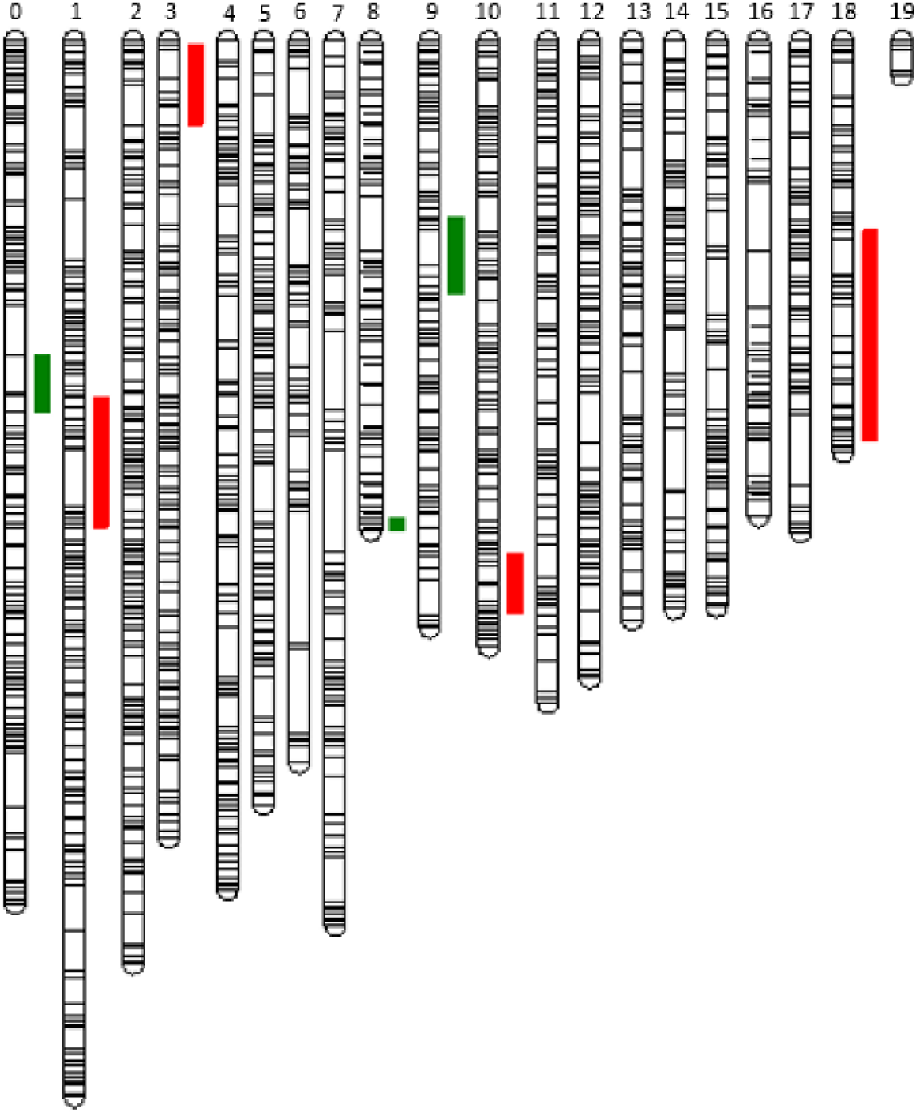
QTL responsible for *Leptosphaeria maculans* pathogenicity on *Brassica napus* (red) and *Brassica carinata* (green) mapped on the genetic map of the cross between JN3 and HB10.19. The numbers represent the linkage groups (LG) of the genetic map and correspond to the chromosome numbers (i.e. Super-Contigs). Each bar line within the LG represents the position of a bin (one skeleton marker associated with its related markers). The visualization of the genetic map was generated using MapChart.

## References

Amezrou, R., Audéon, C., Compain, J., Gélisse, S., Ducasse, A., Saintenac, C., Lapalu, N., Louet, C., Orford, S., Croll, D., Amselem, J., Fillinger, S., Marcel, T.C., 2023, A secreted protease-like protein in *Zymoseptoria tritici* is responsible for avirulence on *Stb9* resistance gene in wheat. Plos Pathog 19: e1011376. 10.1371/journal.ppat.1011376

Amezrou, R., Ducasse, A., Compain, J., Lapalu, N., Pitarch, A., Dupont, L., Confais, J., Goyeau, H., Kema, G.H.J., Croll, D., Amselem, J., Sanchez-Vallet, A., Marcel, T.C., 2024, Quantitative pathogenicity and host adaptation in a fungal plant pathogen revealed by whole-genome sequencing. Nat Commun 15: 1933. 10.1038/s41467-024-46191-1

Ansan-Melayah, D., Balesdent, M.H., Buée, M., Rouxel, T., 1995, Genetic characterization of *AvrLm1*, the first avirulence gene of *Leptosphaeria maculans*. Phytopathology 85: 1525–1529. 10.1094/Phyto-85-1525

Arends, D., Prins, P., Jansen, R.C., Broman, K.W,,2010, R/qtl: high-throughput multiple QTL mapping. Bioinformatics. 26: 2990–2992. https://doi/10.1093/bioinformatics/btq565.

Ayliffe, M., Sørensen, C.K., 2019, Plant nonhost resistance: paradigms and new environments. Current Opinion in Plant Biology 50: 104–113. 10.1016/j.pbi.2019.03.011

Baker, C.J., Harrington, T.C, Krauss, U., Alfena, A.C, 2003, Genetic variability and host specialization in the latin American clade of *Ceratocystis fimbriata*. Phytopathology 93: 1274–1284. 10.1094/PHYTO.2003.93.10.1274

Balesdent, M.H., Fudal, I., Ollivier, B., Bally, P., Grandaubert, J., Eber, F., Chèvre, A.M., Leflon, M., Rouxel, T., 2013, The dispensable chromosome of *Leptosphaeria maculans* shelters an effector gene conferring avirulence towards *Brassica rapa*. New Phytol 198: 887–898. 10.1111/nph.12178

Balesdent, M.H., Attard, A., Ansan-Melayah, D., Delourme, R., Renard, M., Rouxel, T., 2001, Genetic control and host range of avirulence toward *Brassica napus* cultivars Quinta and Jet Neuf in *Leptosphaeria maculans*. Phytopathology 91: 70–76. 10.1094/PHYTO.2001.91.1.70

Balesdent, M.H., Gautier, A., Plissonneau, C., Le Meur, L., Loiseau, A., Leflon, M., Carpezat, J., Pinochet, X., Rouxel, T., 2022, Twenty years of *Leptosphaeria maculans* population survey in France suggests pyramiding *Rlm3* and *Rlm7* in rapeseed is a risky resistance management strategy. Phytopathology 112: 2126–2137. 10.1094/PHYTO-04-22-0108-R

Balint-Kurti, P., 2019, The plant hypersensitive response: concepts, control and consequences. Mol Plant Pathol 20: 1163–1178. 10.1111/mpp.12821

Brown, J.K.M., 2015, Durable resistance of crops to disease: a Darwinian perspective. Annu Rev Phytopathol 53: 513–539. 10.1146/annurev-phyto-102313-045914

Broman, K.W., Wu, H., Sen, S., Churchill, G.A., 2003, R/qtl: QTL mapping in experimental crosses. Bioinformatics 19: 889–890. 10.1093/bioinformatics/btg112.

Caffier, V., Shiller, J., Bellanger, M.N., Collemare, J., Expert, P., Gladieux, P, Pascouau, C., Sannier, M. ^1^, Le Cam, B., 2022, Hybridizations between *formae speciales* of *Venturia inaequalis* pave the way for a new biocontrol strategy to manage fungal plant pathogens. Phytopathology 112: 1401–1405. 10.1094/PHYTO-05-21-0222-SC

Chung, K.R., Hollin, W., Siegel, M.R., Schardl C.L., 1997, Genetics of host specificity in *Epichloë typhina*. Phytopathology 87: 599–605. 10.1094/PHYTO.1997.87.6.599

Cingolani, P., Platts, A., Wang, L.L., Coon, M., Nguyen, T., Wang, L., Land, S.J., Lu, X., Ruden, D.M., 2012, A program for annotating and predicting the effects of single nucleotide polymorphisms, SnpEff: SNPs in the genome of Drosophila melanogaster strain w^1118^ ; *iso*-2; *iso*-3. Fly 6: 80–92. 10.4161/fly.19695

Collemare, J., O’Connell, R., Lebrun, M.H., 2019, Nonproteinaceous effectors: the terra incognita of plant–fungal interactions. New Phytologist 223: 590–596. 10.1111/nph.15785

Couch, B.C., Fudal, I., Lebrun, M.H., Tharreau, D., Valent, B., van Kim, P., Nottéghem, J.L., Kohn, L.M., 2005, Origins of host-specific populations of the blast pathogen *Magnaporthe oryzae* in crop domestication with subsequent expansion of pandemic clones on rice and weeds of rice. Genetics 170: 613–630. 10.1534/genetics.105.041780

Cozijnsen, A.J., Popa, K.M., Purwantara, A., Rolls, B.D., Howlett, B.J., 2000, Genome analysis of the plant pathogenic ascomycete *Leptosphaeria maculans*; mapping mating type and host specificity loci. Mol Plant Pathol 1: 293–302. 10.1046/j.1364-3703.2000.00033.x

Croll, D., Lendenmann, M.H, Stewart, E., McDonald, B.A, 2015, The impact of recombination hotspots on genome evolution of a fungal plant pathogen. Genetics 201: 1213–1228. 10.1534/genetics.115.180968

Cumagun, C. J.R., Rabenstein, F., Miedaner, T., 2004b, Genetic variation and covariation for aggressiveness, deoxynivalenol production and fungal colonization among progeny of *Gibberella zeae* in wheat. Plant Pathol 53: 446–453. 10.1111/j.1365-3059.2004.01046.x

Dahanayaka, B.A., Martin, A., 2023, Multi-parental fungal mapping population study to detect genomic regions associated with *Pyrenophora teres* f. *teres* virulence. Sci Rep 13: 9804. 10.1038/s41598-023-36963-y

Daverdin, G., Rouxel, T., Gout, L., Aubertot, J.N., Fudal, I., Meyer, M., Parlange, F., Carpezat, J., Balesdent, M.H, 2012, Genome structure and reproductive behaviour influence the evolutionary potential of a fungal phytopathogen. PLoS Pathog 8: e1003020. 10.1371/journal.ppat.1003020

Dean, R., Van Kan, J.A.L., Pretorius, Z.A., Hammond-Kosack, K.E., Di Pietro, A., Spanu, P.D., Rudd, J.J., Dickman, M., Kahmann, R., Ellis, J., Foster, G.D., 2012, The Top 10 fungal pathogens in molecular plant pathology. Mol Plant Pathol 13: 414–430. 10.1111/j.1364-3703.2011.00783.x

Delourme, R., Chevre, A.M., Brun, H., Rouxel, T., Balesdent, M.H., Dias, J.S., Salisbury, P., Renard, M., Rimmer, S.R., 2006, Major gene and polygenic resistance to *Leptosphaeria maculans* in oilseed rape (*Brassica napus*). Eur J Plant Pathol 114: 41–52. 10.1007/s10658-005-2108-9

Dutreux, F., Da Silva, C., d’Adata, L., Couloux, A., Gay, E.J., Istace, B., Lapalu, N., Lemainque, A., Linglin, J., Noel, B., Wincker, P., Cruaud, C., Rouxel, T., Balesdent, M.H., Aury, J.M., 2018, De novo assembly and annotation of three *Leptosphaeria* genomes using Oxford Nanopore MinION sequencing. Sci Data 5: 180235. 10.1038/sdata.2018.235

Faino, L., Seidl, M.F., Shi-Kunne, X., Pauper, M., van den Berg, G.C.M., Wittenberg, A.H.J., Thomma, B.P.H.J., 2016, Transposons passively and actively contribute to evolution of the two-speed genome of a fungal pathogen. Genome Res 26: 1091–1100. 10.1101/gr.204974.116

Falconer, D.S., Mackay, T.F.C., 1996, Introduction to Quantitative Genetics. 4th Edition, Addison Wesley Longman, Harlow.

Fisher, M.C., Henk, D.A., Briggs, C.J, Brownstein, J.S., Madoff, L.C., McCraw, S.L., Gurr, S.J., 2012, Emerging fungal threats to animal, plant and ecosystem health. Nature 484: 186–194. 10.1038/nature10947

Fitt, B.D.L., Hu, B.C., Li, Z.Q., Liu, S.Y., Lange, R.M., Kharbanda, P.D., Butterworth, M.H., White, R.P., 2008, Strategies to prevent spread of *Leptosphaeria maculans* (phoma stem canker) onto oilseed rape crops in China; costs and benefits. Plant Pathol 57: 652–664. 10.1111/j.1365-3059.2008.01841.x

Flor, H.H., 1955, Host-parasite interaction in flax-rust. Phytopathology 45: 680–685.

Fokkens, L., Shahi, S., Connolly, L.R., Stam, R., Schmidt, S.M., Smith, K.M., Freitag, M., Martijn Rep, M., 2018, The multi-speed genome of *Fusarium oxysporum* reveals association of histone modifications with sequence divergence and footprints of past horizontal chromosome transfer events. bioRxiv. 10.1101/465070

Fones, H.N., Bebber, D.P., Chaloner, T.M., Kay, W.T., Steinberg, G., Gurr, S.J., 2020, Threats to global food security from emerging fungal and oomycete crop pathogens. Nat Food. 10.1038/s43016-020-0075-0

Fonseca, J.P., Mysore, K.S., 2019, Genes involved in nonhost disease resistance as a key to engineer durable resistance in crops. Plant Science 279: 108–116. 10.1016/j.plantsci.2018.07.002

Fourie, A. Wingfield, M.J., Wingfield, B.D., van der Nest, M.A., Loots, M.T., Barnes, I., 2018, Inheritance of phenotypic traits in the progeny of a *Ceratocystis* interspecific cross. Fungal Biol 122: 717–729. 10.1016/j.funbio.2018.03.001

Fourie, A., van der Nest, M.A., de Vos, L., Wingfield, M.J., Wingfield, B.D., Barnes, I., 2019, QTL mapping of mycelial growth and aggressiveness to distinct hosts in *Ceratocystis* pathogens. Fungal Genet Biol 131: 103242. 10.1016/j.fgb.2019.103242

Fudal, I., Ross, S., Brun, H., Besnard, A.L., Ermel, M., Kuhn, M.L., Balesdent, M.H., Rouxel, T., 2009, Repeat-Induced Point Mutation (RIP) as an alternative mechanism of evolution toward virulence in *Leptosphaeria maculans*. MPMI 22: 932–941. 10.1094/MPMI-22-8-0932

Gay, E.J., Soyer, J.L., Lapalu, N., Linglin, J., Fudal, I., Da Silva, C., Winckler, P., Aury, J.M., Cruaud, C., Levrel, A., Lemoine, J., Delourme, R., Rouxel, T., Balesdent, M.H., 2021, Large-scale transcriptomics to dissect 2 years of the life of a fungal phytopathogen interacting with its host plant. BMC Biol 19: 55. 10.1186/s12915-021-00989-3

Gay, E.J., Jacques, N., Lapalu, N., Cruaud, C., Laval, V., Balesdent, M.H., Rouxel, T., 2023, Location and timing govern tripartite interactions of fungal phytopathogens and host in the stem canker species complex. BMC Biol 21: 247. 10.1186/s12915-023-01726-8

Ghanbarnia, K., Lydiate, D.J., Rimmer, S.R., Li, G., Kutcher, H.R., Larkan, N.J., McVetty, P.B.E., Fernando, W.G.D., 2012, Genetic mapping of the *Leptosphaeria maculans* avirulence gene corresponding to the *LepR1* resistance gene of *Brassica napus*. Theor Appl Genet 124: 505–513. 10.1007/s00122-011-1724-3

Gibson, A.K., Refrégier, G., Hood, M.E., Giraud, T., 2014, Performance of a hybrid fungal pathogen on pure-species and hybrid host plants. Int J Plant Sci 175: 724–730. 10.1086/676621

Gohari M.A., Ware, S.B., Wittenberg, A.H.J., Mehrabi, R., Ben M’Barek, S., Verstappen, E.C.P., van der Lee, T.A.J., Robert, O., Schouten, H.J., de Wit, P.P.J.G.M, Kema, G.H.J., 2015, Effector discovery in the fungal wheat pathogen *Zymoseptoria tritici*. Mol Plant Pathol 16: 931–45. 10.1111/mpp.12251

Grandaubert, J., Lowe, R.G.T., Soyer, J.L., Schoch, C.L., Van de Wouw, A.P., Fudal, I., Robbertse, B., Lapalu, N., Links, M.G., Ollivier, B., Linglin, J., Barbe, V., Mangenot, S., Cruaud, C., Borhan, H., Howlett, B.J., Balesdent, M.H., Rouxel, T., 2014, Transposable element-assisted evolution and adaptation to host plant within the *Leptosphaeria maculans-Leptosphaeria biglobosa* species complex of fungal pathogens. BMC Genomics 15: 891. 10.1186/1471-2164-15-891

Hartmann, F.E., Croll, D., 2017, Distinct Trajectories of massive recent gene gains and losses in populations of a microbial eukaryotic pathogen. Mol Biol Evol 34: 2808–2822. 10.1093/molbev/msx208

Hartmann, F.E., Vonlanthen, T., Singh, N.K., McDonald, M.C., Milgate, A., Croll, D., 2021a, The complex genomic basis of rapid convergent adaptation to pesticides across continents in a fungal plant pathogen. Mol Ecol 30: 5390–5405. 10.1111/mec.15737

Hartmann, F.E., Duhamel, M., Carpentier, F., Hood, M.E., Foulongne-Oriol, M., Silar, P., Malagnac, F., Grognet, P., Giraud, T., 2021b, Recombination suppression and evolutionary strata around mating-type loci in fungi: documenting patterns and understanding evolutionary and mechanistic causes. New Phytol 229: 2470–2491. 10.1111/nph.17039

Jacques, N., Balesdent, M.H., Rouxel, T., Laval V., 2021, New specific quantitative real-time PCR assays shed light on the epidemiology of two species of the *Leptosphaeria maculans* – *Leptosphaeria biglobosa* species complex. Plant Pathol 70: 643–654. 10.1111/ppa.13323

Jiquel, A., Gervais, J., Geistodt-Kiener, A., Delourme, R., Gay, E.J., Ollivier, B., Fudal, I., Faure, S., Balesdent, M.H., Rouxel, T., 2021, A gene-for-gene interaction involving a ‘late’ effector contributes to quantitative resistance to the stem canker disease in *Brassica napus*. New Phytol 231: 1510–1524. 10.1111/nph.17292

Kent, T.V., Uzunović, J., Wright, S.I., 2017, Coevolution between transposable elements and recombination. Phil Trans R Soc B 372: 20160458. 10.1098/rstb.2016.0458

Komluski, J., Stukenbrock, E.H., Habig, M., 2022, Non-Mendelian transmission of accessory chromosomes in fungi. Chromosome Res 30: 241–253. 10.1007/s10577-022-09691-8

Kuhn, M.L., Gout, L., Howlett, B.J., Melayah, D., Meyer, M., Balesdent, M.H., Rouxel, T., 2006, Genetic linkage maps and genomic organization in *Leptosphaeria maculans*. Eur J Plant Pathol 114: 17–31. 10.1007/s10658-005-3168-6

Langlands-Perry, C., Cuenin, M., Bergez, C., Ben Krima, S., Gélisse, S., Sourdille, P., Valade, R., Marcel, T.C., 2021, Resistance of the wheat cultivar ’Renan’ to septoria leaf blotch explained by a combination of strain specific and strain non-specific QTL mapped on an ultra-dense genetic map. Genes 13:100. 10.3390/genes13010100.

Langlands-Perry, C., Pitarch, A., Lapalu, N., Cuenin, M., Bergez, C., Noly, A., Amezrou, R., Gélisse, S., Barrachina, C., Parrinello, H., Suffert, F., Valade, R., Marcel, T.C., 2023, Quantitative and qualitative plant-pathogen interactions call upon similar pathogenicity genes with a spectrum of effects. Front Plant Sci 14: 1128546. 10.3389/fpls.2023.1128546

Laurent, B., Palaiokostas, C., Spataro, C., Moinard, M., Zehraoui, E., Houston, R.D., Foulongne-Oriol, M., 2018, High-resolution mapping of the recombination landscape of the phytopathogen *Fusarium graminearum* suggests two-speed genome evolution. Mol Plant Pathol 19: 341–354. 10.1111/mpp.12524

Laurent, B., Moinard, M., Spataro, C., Chéreau, S., Zehraoui, E., Blanc, R., Lasserre, P., Ponts, N., Foulongne-Oriol, M., 2021, QTL mapping in *Fusarium graminearum* identified an allele of *FgVe1* involved in reduced aggressiveness. Fungal Genet and Biol 153: 103566. 10.1016/j.fgb.2021.103566

Leclair, S., Ansan-Melayah, D., Rouxel, T., Balesdent, M.H., 1996, Meiotic behaviour of the minichromosome in the phytopathogenic ascomycete *Leptosphaeria maculans*. Curr Genet 30: 541–548. 10.1007/s002940050167

Lendenmann, M.H., Croll, D., Stewart, E.L., McDonald, B.A., 2014, Quantitative trait locus mapping of melanization in the plant pathogenic fungus *Zymoseptoria tritici*. G3 4: 2519–2533. 10.1534/g3.114.015289

Lendenmann, M.H., Croll, D., McDonald, B.A., 2015, QTL mapping of fungicide sensitivity reveals novel genes and pleiotropy with melanization in the pathogen *Zymoseptoria tritici*. Fungal Genet and Biol 80: 53–67. 10.1016/j.fgb.2015.05.001

Lendenmann, M.H., Croll, D., Palma-Guerrero, J., Stewart, E.L., McDonald, B.A., 2016, QTL mapping of temperature sensitivity reveals candidate genes for thermal adaptation and growth morphology in the plant pathogenic fungus *Zymoseptoria tritici*. Heredity 116: 384–394. 10.1038/hdy.2015.111

Lo Presti, L., Lanver, D., Schweizer, G., Tanaka, S., Liang, L., Tollot, M., Zuccaro, A., Reissmann, S., Kahmann, R., 2015, Fungal effectors and plant susceptibility. Annu Rev Plant Biol 66: 513–545. 10.1146/annurev-arplant-043014-114623

Louet, C., Saubin, M., Andrieux, A., Persoons, A., Gorse, M., Pétrowski, J., Fabre, B., De Mita, S., Duplessis, S., Frey, P., Halkett, F., 2021, A point mutation and large deletion at the candidate avirulence locus *AvrMlp7* in the poplar rust fungus correlate with poplar RMlp7 resistance breakdown. Mol Ecol 32: 2472–2483 10.1111/mec.16294

McDonald, B.A., Linde, C., 2002, Pathogen population genetics, evolutionary potential, and durable resistance. Annu Rev Phytopathol 40: 349–379. 10.1146/annurev.phyto.40.120501.101443

Meile, L., Croll, D., Brunner, P.C., Plissonneau, C., Hartmann, F.E., McDonald, B.A., Sánchez-Vallet, A., 2018, A fungal avirulence factor encoded in a highly plastic genomic region triggers partial resistance to septoria tritici blotch. New Phytol 219: 1048–1061. 10.1111/nph.15180

Meile, L., Garrido-Arandia, M., Bernasconi, Z., Peter, J., Schneller, A., Bernasconi, A., Alassimone, J., McDonald, B.A., Sánchez-Vallet, A., 2023, Natural variation in *Avr3D1* from *Zymoseptoria* sp. contributes to quantitative gene-for-gene resistance and to host specificity. New Phytol 238: 1562–1577. 10.1111/nph.18690

Mester, D., Ronin, Y., Minkov, D., Nevo, E., Korol, A., 2003, Constructing large-scale genetic maps using an evolutionary strategy algorithm. Genetics 165: 2269–2282.

Mézard, C., Tagliaro Jahns, M., Grelon, M., 2015, Where to cross? New insights into the location of meiotic crossovers. Trends Genet 31: 393–401. 10.1016/j.tig.2015.03.008

Miñana-Posada, S., Lorrain, C., McDonald, B.A., Feurtey, A., 2025, Thermal adaptation in worldwide collections of a major fungal pathogen. MPMI 38: 252–264. 10.1094/MPMI-09-24-0112-FI

Nelson, R., Wiesner-Hanks, T., Wisser, R., Balint-Kurti, P., 2018, Navigating complexity to breed disease-resistant crops. Nat Rev Genet 19: 21–33. 10.1038/nrg.2017.82

Niks, R.E., Marcel, T.C., 2009, Nonhost and basal resistance: how to explain specificity? New Phytol 182: 817–828. 10.1111/j.1469-8137.2009.02849.x

Noah, J.M., Gorse, M., Romain, C.A., Gay, E.J., Rouxel, T., Balesdent, M.H., Soyer, J.L., 2024a, To be or not to be a nonhost species: A case study of the *Leptosphaeria maculans* and *Brassica carinata* interaction. Environ Microbiol Rep 16: e70034. 10.1111/1758-2229.70034

Noah, J.M., Lapalu, N., Rouxel, T., Balesdent, M.H., Soyer, J.L., 2024b, *Leptosphaeria maculans* semi-quantitative disease scale (LeMaSQ). Recherche Data Gouv, V1 10.57745/OYUO2X

Panstruga, R., Moscou, M.J., 2020, What is the molecular basis of nonhost resistance? MPMI 33: 1253–1264. 10.1094/MPMI-06-20-0161-CR

Pongam, P., Osborn, T.C., Williams, P.H., 1998, Assessment of genetic variation among *Leptosphaeria maculans* isolates using pathogenicity data and AFLP analysis. Plant Disease 83: 149–154. 10.1094/PDIS.1999.83.2.149

Rieseberg, L.H., Archer, M.A., Wayne, R.K., 1999, Transgressive segregation, adaptation and speciation. Heredity 83: 363–372. 10.1038/sj.hdy.6886170

Rieseberg, L.H., Widmer, A., Arntz, A.M., Burke, J.M., 2003, The genetic architecture necessary for transgressive segregation is common in both natural and domesticated populations. Phil Trans R Soc Lond 358: 1141–1147. 10.1098/rstb.2003.1283

Ronin, Y.I., Mester, D.I., Minkov, D.G., Akhunov, E., Korol, A.B., 2017, Building ultra-high-density linkage maps based on efficient filtering of trustable markers. Genetics 206: 1285–1295. 10.1534/genetics.116.197491

Rouxel, T., Penaud, A., Pinochet, X., Brun, H., Gout, L., Delourme, R., Schmit, J., Balesdent, M.H., 2003, A 10-year survey of populations of *Leptosphaeria maculans* in France indicates a rapid adaptation towards the *Rlm1* resistance gene of oilseed rape. Eur J Plant Pathol 109: 871–881. 10.1023/A:1026189225466

Rouxel, T., Balesdent, M.H., 2005, The stem canker (blackleg) fungus, *Leptosphaeria maculans*, enters the genomic era. Mol Plant Pathol 6: 225–241. 10.1111/j.1364-3703.2005.00282.x

Rouxel, T., Balesdent, M.H., 2017, Life, death and rebirth of avirulence effectors in a fungal pathogen of *Brassica* crops, *Leptosphaeria maculans*. New Phytol 214: 526–532. 10.1111/nph.14411

Rouxel, T., Grandaubert, J., Hane, J.H., Hoede, C., van de Wouw, A.P., Couloux, A., Dominguez, V., Anthouard, V., Bally, P., Bourras, S., Cozijnsen, A.J., Ciuffetti, L.M., Degrave, A., Dilmaghani, A., Duret, L., Fudal, I., Goodwin, S.B., Gout, L., Glaser, N., Linglin, J., Kema, G.H.J., Lapalu, N., Lawrence, C.B., May, K., Howlett B.J., 2011, Effector diversification within compartments of the *Leptosphaeria maculans* genome affected by Repeat-Induced Point mutations. Nat Commun 2: 202. 10.1038/ncomms1189

Sánchez-Vallet, A., Fouché, S., Fudal, I., Hartmann, F.E., Soyer, J.L., Tellier, A., Croll, D., 2018, The genome biology of effector gene evolution in filamentous plant pathogens. Annu Rev Phytopathol 56: 21–40. 10.1146/annurev-phyto-080516-035303

Schneider, C.A., Rasband, W.S., Eliceiri, K.W., 2012, NIH Image to ImageJ: 25 years of image analysis. Nat Methods 9: 671–675. 10.1038/nmeth.2089

Schulze-Lefert, P., Panstruga, R., 2011, A molecular evolutionary concept connecting nonhost resistance, pathogen host range, and pathogen speciation. Trends in Plant Science 16: 117–125. 10.1016/j.tplants.2011.01.001

Seidl, M.F., Thomma, B.P.H.J., 2017, Transposable elements direct the coevolution between plants and microbes. Trends Genet 33: 842–851. 10.1016/j.tig.2017.07.003

Sharma, R., Mishra, B., Runge, F., Thines M., 2014, Gene loss rather than gene gain is associated with a host jump from monocots to dicots in the smut fungus *Melanopsichium pennsylvanicum*. Genome Biol Evol 6: 2034–2049. 10.1093/gbe/evu148

Sonnenberg, A.S.M., Baars, J.J.P., Gao, W., Visser, R.G.F., 2017, Developments in breeding of *Agaricus bisporus var. bisporus*: progress made and technical and legal hurdles to take. Appl Microbiol Biotechnol 101: 1819–1829. 10.1007/s00253-017-8102-2

Soyer, J.L., El Ghalid, M., Glaser, N., Ollivier, B., Linglin, J., Grandaubert, J., Balesdent, M.H., Connolly, L.R., Freitag, M., Rouxel, T., Fudal, I., 2014, Epigenetic control of effector gene expression in the plant pathogenic fungus *Leptosphaeria maculans*. Plos Genet 10: e1004227. 10.1371/journal.pgen.1004227

Soyer, J.L., Clairet, C., Gay, E.J., Lapalu, N., Rouxel, T., Stukenbrock, E.H., Fudal, I., 2021, Genome-wide mapping of histone modifications during axenic growth in two species of *Leptosphaeria maculans* showing contrasting genomic organization. Chromosome Res 29: 219–236. 10.1007/s10577-021-09658-1

Stapley, J., Zhong, Z., McDonald, B.A., 2025, Mapping genomic regions associated with temperature stress in the wheat pathogen *Zymoseptoria tritici*. G3 15. 10.1093/g3journal/jkaf094

Stelkens, R., Seehausen, O., 2009, Genetic distance between species predicts novel trait expression in their hybrids. Evolution 63: 884–897. 10.1111/j.1420-9101.2009.01777.x

Stewart, E.L., Croll, D., Lendenmann, M.H., Sanchez-Vallet, A., Hartmann F.E., Palma-Guerrero, J., Ma, X., McDonald, B.A., 2018, Quantitative trait locus mapping reveals complex genetic architecture of quantitative virulence in the wheat pathogen *Zymoseptoria tritici*: QTL mapping of virulence in *Zymoseptoria tritici*. Molecular Plant Pathology 19: 201–216. 10.1111/mpp.12515

Tarigan, M., Roux, J., Van Wyk, M., Tjahjono, B., Wingfield, M.J., 2011, A new wilt and die-back disease of *Acacia mangium* associated with *Ceratocystis manginecans* and *C. acaciivora sp. nov.* in Indonesia. S Afr J Bot 77: 292–304. 10.1016/j.sajb.2010.08.006

Thines, M., 2019, An evolutionary framework for host shifts – jumping ships for survival. New Phytol 224: 605–617. 10.1111/nph.16092

Torres, D.E., Oggenfuss, U, Croll, D., Seidl, M.F., 2020, Genome evolution in fungal plant pathogens: looking beyond the two-speed genome model. Fungal Biol Rev 34: 136–143. 10.1016/j.fbr.2020.07.001

Valdetaro, D.C.O.F., Harrington, T.C., Oliveira, L.S.S., Guimaraes, L.M.S., McNew, D.L., Pimenta, L.V.A., Gonçalves, R.C., Schurt, D.A., Alfenas, A.C., 2019, A host specialized form of *Ceratocystis fimbriata* causes seed and seedling blight on native *Carapa guianensis* (andiroba) in Amazonian rainforests. Fungal Biol 123: 170–182. 10.1016/j.funbio.2018.12.001

van Dam, P., Fokkens, L., Schmidt, S.M., Linmans, J.H.J., Kistler, H.C., Ma, L.J., Rep, M., 2016, Effector profiles distinguish *formae speciales* of *Fusarium oxysporum*. Environ Microbio 18: 4087–4102. doi: 10.1111/1462-2920.13445

Vasquez-Teuber, P., Rouxel, T., Mason, A.S., Soyer, J.L., 2024, Breeding and management of major resistance genes to stem canker/blackleg in *Brassica* crops. Theor Appl Genet 137: 192. 10.1007/s00122-024-04641-w

Voorrips, R.E., 2002, MapChart: software for the graphical presentation of linkage maps and QTLs. J Hered 93: 77–78. 10.1093/jhered/93.1.77

Voss, H.H., Bowden, R.L., Leslie, J.F., Miedaner, T., 2010, Variation and transgression of aggressiveness among two *Gibberella zeae* crosses developed from highly aggressive parental isolates. Phytopathology 100: 904–912. 10.1094/PHYTO-100-9-0904

West, J.S., Kharbanda, P.D., Barbetti, M.J., Fitt, B.D.L., 2001, Epidemiology and management of *Leptosphaeria maculans* (phoma stem canker) on oilseed rape in Australia, Canada and Europe: epidemiology of L. maculans on rapeseed. Plant Pathol 50: 10–27. 10.1046/j.1365-3059.2001.00546.x

Zhong, Z., Marcel, T.C., Hartmann, F.E., Ma, X., Plissonneau, C., Zala, M., Ducasse, A., Confais, J., Compain, J., Lapalu, N., Amselem, J., McDonald, B.A., Croll, D., Palma-Guerrero, J., 2017, A small secreted protein in *Zymoseptoria tritici* is responsible for avirulence on wheat cultivars carrying the *Stb6* resistance gene. New Phytol 214: 619–631. 10.1111/nph.14434

